# Characterization of BioID tagging systems in budding yeast and exploring the interactome of the Ccr4-Not complex

**DOI:** 10.1101/2024.05.09.593354

**Authors:** J. Pfannenstein, M. Tyryshkin, M.E Gulden, E.H. Doud, A. L. Mosley, J.C. Reese

## Abstract

The modified E. coli biotin ligase BirA* was the first developed for proximity labeling of proteins (BioID). However, it has low activity at temperatures below 37°C, which reduces its effectiveness in organisms growing at lower temperatures, such as budding yeast. Multiple derivatives of the enzymes have been engineered, but a comparison of these variations of biotin ligases has not been reported in Saccharomyces cerevisiae. Here, we designed a suite of vectors to compare the activities of biotin ligase enzymes in yeast. We found that the newer TurboID versions were the most effective at labeling proteins, but they displayed low constitutive activity from biotin contained in the culture medium. We describe a simple strategy to express free BioID enzymes in cells that can be used as an appropriate control in BioID studies to account for the promiscuous labeling of proteins caused by random interactions between bait-BioID enzymes in cells. We also describe chemically-induced BioID systems exploiting the rapamycin-stabilized FRB-FKBP interaction. Finally, we used the TurboID version of the enzyme to explore the interactome of different subunits of the Ccr4-Not gene regulatory complex. We find that Ccr4-Not predominantly labeled cytoplasmic mRNA regulators, consistent with its function in mRNA decay and translation quality control in this cell compartment.

## Introduction

Understanding biological processes requires, among other techniques, mapping protein-protein interactions (PPI). This is often performed by affinity capture methods of a bait protein, followed by mass spectrometry to identify the interacting proteins [1]. This approach has generated rich, expansive datasets to complement genetic and molecular analysis of cellular pathways. However, preparing extracts takes proteins out of their cellular context, and the dilution of components during purification and the stringent conditions required to reduce background can disrupt biologically relevant PPIs. Furthermore, these methods capture stable, long-lived interactions but often fail to identify transient interactions. A few procedures have been developed to alleviate these shortcomings. One such technique is proximity labeling of proteins in live cells using engineered *E. coli* biotin ligases [1, 2]. The modified enzyme is fused in-frame with a protein of interest, or bait protein. After the addition of biotin, proteins within a certain radius of the enzyme, estimated to be 10nM, are biotinylated on accessible lysine residues. The biotin-streptavidin interaction’s unusual strength allows for capturing the biotinylated proteins under conditions that break non-covalent PPIs and reduce background binding.

The first enzyme developed for this application contained a R118G mutation in *E. Coli* BirA, referred to as BirA*[1, 2]. This mutation reduces the affinity of the bioAMP product, which diffuses from the enzyme and covalently attaches to lysine residues of adjacent proteins. Another version, termed BioID2, reduced the size of the enzyme and increased its activity [3]. One drawback of the earlier BirA* versions is that they display slow kinetics, requiring many hours of labeling (16-24hrs), and low activity below 37°C [4]. Both features limit their use in organisms that grow below 37°C and display short cell cycles, such as budding yeast. However, BirA* has been used in yeast to a limited degree on abundant proteins [5]. A version of biotin ligase from Bacillus subtilis, termed BASU, was engineered that appears to be more active than BirA*. Since BASU originated from a microbe that grows at a lower temperature than E. coli, it might be more amenable to use in a wider variety of organisms To our knowledge, BASU has not been tested in yeast. A significant advance in BioID studies was made through the directed evolution and selection of a new biotin ligase termed TurboID, TID [4]. TID was selected in a yeast surface display system conducted at lower temperatures, which resulted in very high activity at lower temperatures. A shortened version, miniTurbo ID (mTID), was identified and is about 2-fold less active. A comparison of the new TID versions versus the original BirA* demonstrated it was much more active [4]. A potential drawback of the TID and mTID enzymes is that they biotinylated proteins without adding exogenous biotin to the media [4, 6]. Many BioID studies rely upon comparing captured proteins plus and minus biotin, and the constitutive biotinylation from TID complicates this strategy. Furthermore, the expression of biotin ligase fusion proteins can lead to spurious biotinylation of proteins due to random collisions with cellular proteins, which is particularly problematic for abundant proteins.

The TurboID version has been used in yeast to define the interactome of the protein arginine methyltransferase Rmt3 and the exosome subunits Dis3 and Rrp6 [7]. This study designed a limited vector system to tag proteins with TID [7]. However, the yeast field lacks a versatile vector system to tag proteins with different versions of biotin ligases. Moreover, a direct and comprehensive comparison of the collection of BioID enzymes has yet to be performed in budding yeast. Here, we design a modular vector system to tag and analyze the biotin ligases BirA*, BioID2, BASU, TurboID, and miniTurboID in yeast. Furthermore, we designed and tested a chemically induced BioID system that exploits the rapamycin-dependent FRB-FKBP interaction. We describe a simple and rapid method to generate strains expressing “free” TurboID that can be used as a control in BioID studies. We applied the BioID system to identify the interactome of the Ccr4-Not complex.

## Methods

### Yeast strains

Yeast strains are listed in Supplemental Table 1. Standard genetic analysis techniques were applied. Integration of genes was confirmed by PCR and/or western blotting to confirm the addition and expression of the tags.

### Construction of BioID plasmids

Plasmids are described in Table 1 and all plasmids will be submitted to Addgene for distribution. Plasmids to insert biotin ligases at the C-terminus of genes were constructed in pF6a-3HA-KanMX6 and pF6a-3HA-HIS3MX6 [8] by digesting the vectors with PacI and then inserting biotin ligases using inphusion gene assembly (Takara). The fragments used in cloning were generated by gene synthesis, and the codons in coding regions were optimized for yeast. Primers containing 45 nucleotides homologous to the target gene, followed by the F2 and R1 primer sequences complementary to the plasmid, were used to amplify gene cassettes for homologous recombination. [8]. PCR cassettes to integrate free biotin ligases downstream of yeast promoters were generated using the same plasmids, but primers containing 45 nts of homologous sequences in the promoters of genes followed by 18-19 nucleotides homologous to the start codon and ORF of the biotin ligase and the sequences in the 3”UTR of the gene followed the R1 sequence were to amplify the cassette. The plasmids used to integrate biotin ligases-FRB/FKBP fusions at yeast promoters were constructed by placing codon-optimized gene fragments into the PacI and AscI sites of pF6a vectors.

**Table 1.** plasmids.

| # | features | description | ref |
| --- | --- | --- | --- |
|  | pFA6a -HA3-KanMX6 | HA3 tagging | 8 |
|  | pFA6a-HA3-HIS3MX6 | HA3 tagging | 8 |
| pJR100 | pFA6a-BirA-HA3-KanMX6 | BirA tagging | This study |
| pJR101 | pFA6a-BirA-HA3-HIS3MX6 | BirA tagging | This study |
| pJR102 | pFA6a-BioID2-HA3-KanMX6 | BioID2 tagging | This study |
| pJR103 | pFA6a-BioID2-HA3-HIS3MX6 | BioID2 tagging | This study |
| pJR104 | pFA6a-BASU-HA3-KanMX6 | BASU tagging | This study |
| pJR105 | pFA6a-BASU-HA3-HIS3MX6 | BASU tagging | This study |
| pJR106 | pFA6a-TID-HA3-KanMX6 | Turbo ID (TID) tagging | This study |
| pJR107 | pFA6a-TID-HA3-HIS3MX6 | Turbo ID (TID) tagging | This study |
| pJR108 | pFA6a-mTID-HA3-KanMX6 | Mini-Turbo ID (mTID) tagging | This study |
| pJR109 | pFA6a-mTID-HA3-HIS3MX6 | Mini-Turbo ID (mTID) tagging | This study |
| P30578 | pFA6a-FRB-KanMX6 | FRB tagging | Euroscarf |
| P30583 | pFA6a-2xFKBP12-His3MX6 | 2xFKBP tagging | Euroscarf |
| pJR110 | pRS416-ADH1p-BirA-FRB-HA3 | BirA-FRB plasmid | This study |
| pJR111 | pRS416-ADH1p-BirA-FRB-NLS-HA3 | BirA-FRB-NLS plasmid | This study |
| pJR112 | pRS416-ADH1p-BirA-FKBP- HA3 | BirA-FKBP plasmid | This study |
| pJR113 | pRS416-ADH1p-BirA-FKBP- NLS-HA3 | BirA-FKBP-NLS plasmid | This study |
| pJR114 | pRS416-ADH1p-BioID2-FRB-HA3 | BioID2-FRB plasmid | This study |
| pJR115 | pRS416-ADH1p- BioID2-FRB-NLS-HA3 | BioID2-FRB-NLS plasmid | This study |
| pJR116 | pRS416-ADH1p- BioID2-FKBP-HA3 | BioID2-FKBP plasmid | This study |
| pJR117 | pRS416-ADH1p- BioID2-FKBP-NLS-HA3 | BioID2-FKBP-NLS plasmid | This study |
| pJR118 | pRS416-ADH1p-BASU-FRB-HA3 | BASU-FRB plasmid | This study |
| pJR119 | pRS416-ADH1p-BASU-FRB-NLS-HA3 | BASU-FRB-NLS plasmid | This study |
| pJR120 | pRS416-ADH1p-BASU-FKBP- HA3 | BASU-FKBP plasmid | This study |
| pJR121 | pRS416-ADH1p-BASU-FKBP- NLS-HA3 | BASU-FKBP-NLS plasmid | This study |
| pJR122 | pRS416-ADH1p-BASU-FRB-HA3 OPT | BASU-FRB codon optimized | This study |
| pJR123 | pRS416-ADH1p-TID-FRB-HA3 | TID-FRB plasmid | This study |
| pJR124 | pRS416-ADH1p-TID-FRB-NLS-HA3 | TID-FRB-NLS plasmid | This study |
| pJR125 | pRS416-ADH1p-TID-FKBP-HA3 | TID-FKBP plasmid | This study |
| pJR126 | pRS416-ADH1p-TID-FKBP-NLS-HA3 | TID-FKBP-NLS plasmid | This study |
| pJR127 | pRS416-ADH1p-mTID-FRB-HA3 | mTID-FRB plasmid | This study |
| pJR128 | pRS416-ADH1p-mTID-FRB-NLS-HA3 | mTID-FRB-NLS plasmid | This study |
| pJR129 | pRS416-ADH1p-mTID-FKBP-HA3 | mTID-FKBP plasmid | This study |
| pJR130 | pRS416-ADH1p-mTID-FKBP-NLS-HA3 | mTID-FKBP-NLS plasmid | This study |
| pJR131 | pRS416-HIS3p-TID-FRB-HA3 | TID-FRB plasmid | This study |
| pJR132 | pRS416-HIS3p-TID-FRB-NLS-HA3 | TID-FRB-NLS plasmid | This study |
| pJR133 | pRS416-HIS3p-TID-FKBP-HA3 | TID-FKBP plasmid | This study |
| pJR134 | pRS416-HIS3p-TID-FKBP-NLS-HA3 | TID-FKBP-NLS plasmid | This study |
| pJR135 | pRS416-CUP1p-TID-FRB-HA3 | TID-FRB plasmid | This study |
| pJR136 | pRS416-CUP1p-TID-FRB-NLS-HA3 | TID-FRB-NLS plasmid | This study |
| pJR137 | pRS416-CUP1p-TID-FKBP-HA3 | TID-FKBP plasmid | This study |
| pJR138 | pRS416-CUP1p-TID-FKBP-NLS-HA3 | TID-FKBP-NLS plasmid | This study |
| pJR139 | pRS416-CHA1p-TID-FRB-HA3 | TID-FRB plasmid | This study |
| pJR140 | pRS416-CHA1p-TID-FRB-NLS-HA3 | TID-FRB-NLS plasmid | This study |
| pJR141 | pRS416-CHA1p-TID-FKBP-HA3 | TID-FKBP plasmid | This study |
| pJR142 | pRS416-CHA1p-TID-FKBP-NLS-HA3 | TID-FKBP-NLS plasmid | This study |
| pJR143 | pRS416-MET25p-TID-FRB-HA3 | TID-FRB plasmid | This study |
| pJR144 | pRS416-MET25p-TID-FRB-NLS-HA3 | TID-FRB-NLS plasmid | This study |
| pJR145 | pRS416-MET25p-TID-FKBP-HA3 | TID-FKBP plasmid | This study |
| pJR146 | pRS416-MET25p-TID-FKBP-NLS-HA3 | TID-FKBP-NLS plasmid | This study |
| pJR147 | pF6a-TID-FRB-3HA-KanMX6 | TID-FRB integration | This study |
| pJR148 | pF6a-TID-FRB-3HA-HIS3MX6 | TID-FRB integration | This study |
| pJR149 | pF6a-TID-FRB-NLS-3HA-KanMX6 | TID-FRB-NLS integration | This study |
| pJR150 | pF6a-TID-FRB-NLS-3HA-HIS3MX6 | TID-FRB-NLS integration | This study |
| pJR151 | pF6a-TID-FKBP-3HA-KanMX6 | TID-FKBP integration | This study |
| pJR152 | pF6a-TID-FKBP-3HA-HIS3MX6 | TID-FKBP integration | This study |
| pJR153 | pF6a-TID-FKBP-NLS-3HA-KanMX6 | TID-FKBP-NLS integration | This study |
| pJR154 | pF6a-TID-FKBP-3HA-NLS-HIS3MX6 | TID-FKBP-NLS integration | This study |

Versions containing a nuclear localization sequence were also constructed (Table 1). Forward and reverse primers for PCR were similar to those used to prepare cassettes to express free biotin ligases at chromosomal loci. Plasmid-expressed biotin ligase-FRB and -FKBP pairs were constructed in pQWF (provided by James Hopper), which is a derivative of pRS416 containing the ADH1 promoter and terminator. Gene synthesis fragments were inserted into the HindIIII and XbaI sites by inphusion reaction. To construct TID-FRB and TID-FKBP versions driven by the *HIS3*, *CHA1, CUP1*, and *MET25* promoters, the ADH1 promoter was removed by digestion with KpnI and HindIII, and PCR-generated promoter fragments were inserted. Gene synthesis and primer sequences are available upon request. Plasmid sequences will be submitted to Addgene.

### Western and streptavidin blotting

Cell extracts for screening protein expression were prepared from 5 OD600 of cells by glass bead disruption in the presence of 20% trichloroacetic acid (TCA). After adjusting the TCA concentration to 10%, the lysate was separated from the beads and centrifuged at 12,000xg. Pellets were rinsed with 1 ml 0.2M Tris-HCl ph 7.5 and resuspended in 150ul 0.1M Tris-HCl pH 9. After the addition of 80 ul 3X SDS-PAGE loading, the samples were boiled and clarified by centrifugation at 14,000xg at room temperature. Typically, 5 ul was loaded onto gels.

Samples were separated on SDS-PAGE gels or 4-15% Tris-acetate gels. The HA tag was detected using 12CA5 (Novus Biologicals) or HA11.1 (Covance, now Labcorp) monoclonal antibodies. Antibodies to Not1, Not4, Caf1, Dhh1, TAF1 and TAF14 were raised in house in rabbits. Streptavidin blots were performed using Ultra Streptavidin-HRP at 1:20,000 and Pico ECL solution (Thermo Scientific) and images were documented with x-ray film or digitally with a ChemiDoc MP (Biorad). The following fluorescent secondary antibodies were used goat anti-Rabbit CY5 (Invitrogen), goat anti-rabbit Starbright Blue 520 and 700 (Biorad) and goat anti-mouse IR800 (Licor). Images were captured on a Typhoon instrument (CY5) (GE Life Sciences) or ChemiDoc MP (Biorad). Images were quantified using ImageJ.

### Extract preparation and affinity capture

Typically, cells were grown in 50 mls synthetic complete media supplemented with 2% dextrose to an OD600 of 0.5-0.6 and then induced with biotin for 2 hrs unless indicated otherwise. Cells were harvested, washed twice in STE (10 mM Tris-HCL, 50 mM NaCl and 1 mM EDTA) containing 1 mM benzamidine-HCl, and snap frozen and stored at -80°C. Cells were resuspended in 500 ul of lysis buffer (20 mM HEPES-NaOH ph 7.5, 0.4 M NaCl, 2 mM EGTA, 10 mM MgCl2, 2 mM DTT,10% glycerol supplemented with the protease inhibitors 1 mM benzamidine-HCl and 1 ug/ml leupeptin, aprotinin and pepstatin A) and distributed into two screw-top microfuge tubes containing glass beads. PMSF was added to 1 mM immediately before lysis. Cells were broken with glass beads using six cycles of 30 seconds on and 30 seconds off at 4°C using a Beadblaster 24 (Benchmark Scientific). Another 250 ul of lysis buffer was added to each tube, and fresh PMSF was added to 1 mM. The glass beads were separated from the lysate, and benzonase (Millipore), and Triton-X-100 was added to 500U/ml and 0.1%, respectively. The cell lysate was incubated at 4°C with rotation for 20-25 minutes.

The lysate was clarified by three cycles of centrifugation at 4°C in a microfuge at 13.5K rpm. Protein concentration was estimated by Bradford assay using BSA as a standard. To enrich proteins for BioID, 1.5 milligrams of protein was diluted up to 1.5 mls in lysis buffer +0.1% triton X-100 and 60 ul of streptavidin magnetic beads (Pierce,#.88817) was added and incubated 16-17 hrs at 4°C with rotation. Beads were recovered on a magnet and washed as follows: three times for 10 mins with lysis buffer+0.1% triton X-100 plus protease inhibitors, once 10 minutes with 10 mM Tris-HCl ph 8, 2 M Urea, 0.3% SDS, once 10 min in FA-lysis buffer 2 (50 mM HEPES/KOH pH 7.5, 2 mM EDTA, 1% Triton X-100, 0.1% sodium deoxycholate; 0.5 M NaCl, 0.5 M PMSF), once for 10 minutes FA-lysis buffer 3 (10 mM Tris-HCl ph 8, 0.25 M LiCl; 1% NP-40; 1% sodium deoxycholate; 2 mM EDTA). Beads were resuspended in STE and transferred to a new tube. The beads were washed once more in STE and then frozen at -80°C until digestion.

One milligram of protein was diluted in lysis buffer containing 0.1% NP40 up to 1.2 mls for immunoprecipitation experiments. Four microliters of pre-immune or anti- Not4 antiserum were added and incubated for 2-3 hrs on ice. The extract was clarified by a 5 min centrifugation step at high speed and transferred to a new tube.

Fifty microliters of a 50-50 slurry of protein A-agarose beads (UBPbio) in lysis buffer was added, and the tubes were incubated overnight at 4°C. After recovery, the beads were washed 4X with lysis buffer containing 0.1% NP40 for 10 minutes each. The proteins were eluted in 40ul of 1.5X SDS-PAGE loading buffer and separated on SDS-PAGE gels or 4-15% Tris-acetate gradient gels where indicated.

### Protein digestion and mass spectrometry

On bead samples were submitted to the IUSM Center for proteome analysis where proteins were denatured in 8 M urea, 100 mM Tris-HCl, pH 8.5 and reduced with 5 mM tris(2-carboxyethyl)phosphine hydrochloride (TCEP, Sigma-Aldrich Cat No: C4706) for 30 minutes at room temperature. Samples were then alkylated with 10 mM chloroacetamide (CAA, Sigma Aldrich Cat No: C0267) for 30 min at room temperature in the dark, prior to dilution with 50 mM Tris. HCl, pH 8.5 to a final urea concentration of 2 M for Trypsin/Lys-C based overnight protein digestion at 37 °C (0.5 µg protease, Mass Spectrometry grade, Promega Corporation, Cat No: V5072.)

Digestions were acidified with trifluoroacetic acid (TFA, 0.5% v/v) and desalted on Pierce C18 spin columns (Thermo Fisher Cat No: 89870) with a wash of 0.5% TFA followed by elution in 70% acetonitrile 0.1% formic acid (FA). Peptides were dried by speed vacuum and resuspended in 24 µL of 50 mM triethylammonium bicarbonate (TEAB). Samples were then Tandem Mass Tag (TMT) labeled with 0.2 mg of reagent according to manufacturers instructions (Thermo Fisher Cat No: A37725. Labelling reactions were quenched with hydroxylamine at room temperature 15 minutes. Labelled peptides were then mixed by global and dried by speed vacuum.

Mass spectrometry was performed utilizing an EASY-nLC 1200 HPLC system (SCR: 014993, Thermo Fisher Scientific) coupled to Lumos™ Tribrid orbitrap (Thermo Fisher Scientific). 1/3^rd^ of the sample was pressure bomb loaded onto a homemade MudPIT column of 20 cm reverse phase resin and 5 cm strong cation exchange resin column and eluted in a 10 steps of increasing ammonium acetate as described previously [9, 10]. The gradient was held at 5% B for 5 minutes (Mobile phases A: 0.1% formic acid (FA), water; B: 0.1% FA, 80% Acetonitrile (Thermo Fisher Scientific Cat No: LS122500)), then increased from 4-30%B over 98 minutes; 30-80% B over 10 mins; held at 80% for 2 minutes; and dropping from 80-4% B over the final 5 min. Mass spectrometer settings include capillary temperature of 300 °C and ion spray voltage was kept at 1.9 kV. The mass spectrometer method was operated in positive ion mode with a 4 second cycle time data-dependent acquisition with advanced peak determination and Easy-IC on (internal calibrant). Precursor scans (m/z 400-1750) were done with an orbitrap resolution of 120000, 30% RF lens, 50 ms maximum inject time (IT) and standard automatic gain control (AGC) target, including charges of 2 to 6 for fragmentation with 45 s dynamic exclusion. Higher-energy collisional dissociation (HCD) MS2 scans were performed at 50k orbitrap resolution, fixed collision energy of 34%, 100% normalized AGC target, and 90 ms maximum IT.

### Mass spectrometry Data Analysis

Resulting RAW files were analyzed in Proteome Discover™ 2.3 or 2.4 (Thermo Fisher Scientific) with a *Saccharomyces cerevisiae* reference proteome FASTA plus common contaminants (73 entries)[11]. SEQUEST HT searches were conducted with a maximum number of 2 missed cleavages; precursor mass tolerance of 10 ppm, and a fragment mass tolerance of 0.6 Da. Static modifications used for the search were carbamidomethylation on cysteine (C), TMT (+229.153) on peptide N-terminus and lysine (K). Dynamic modifications included and oxidation of methionine (M), and acetylation on protein N-termini. Percolator False Discovery Rate was set to a strict peptide spectral match FDR setting of 0.01 and a relaxed setting of 0.05. In the consensus workflow, unique and razor peptides were used for quantification; reporter ion abundance was set to ‘automatic or S/N), Isobaric impurities were not utilized, a co-isolation threshold off 50% and a S/N threshold of 10 were used. Abundances were normalized to total peptide amount in each TMT channel, all peptides were used for normalization and protein-roll-up. Protein ratio calculations were protein abundance based, with no imputation and ANOVA hypothesis testing to calculate p-values.

Raw and searched data will be uploaded to MassIVE repository.

## Results

### Comparing biotin ligase activity in yeast

Multiple promiscuous biotin ligases have been developed since BirA*, each with advantages and disadvantages. We have designed a versatile system to tag genes at the C-terminus with biotin ligases (Figure 1 and Table 1) using the pF6a vector platform[8], which allows for the use of a common set of primers, F2 and R1, and existing vectors that can be used to add different tags (e.g. epitope tags, EGFP). The enzyme sequences were codon-optimized for yeast. To test the activity of the various biotin ligases, we used Not4, a subunit of the Ccr4-Not complex [12–14]. BirA*, BioID2, BASU, TurboID (TID), and mini-TurboID (mTID) were introduced into the C-terminus of Not4 by homologous recombination. First, we confirmed that fusing the biotin ligases to Not4 did not impair gene function. NOT4 mutant cells display slowed growth, temperature- and hydroxyurea-sensitivity [15, 16]. Cells expressing the different Not4-ligase fusions displayed normal growth phenotypes, indicating that fusing the enzyme to Not4 did not impair its function (Supplemental Figure 1).

**Figure 1.**
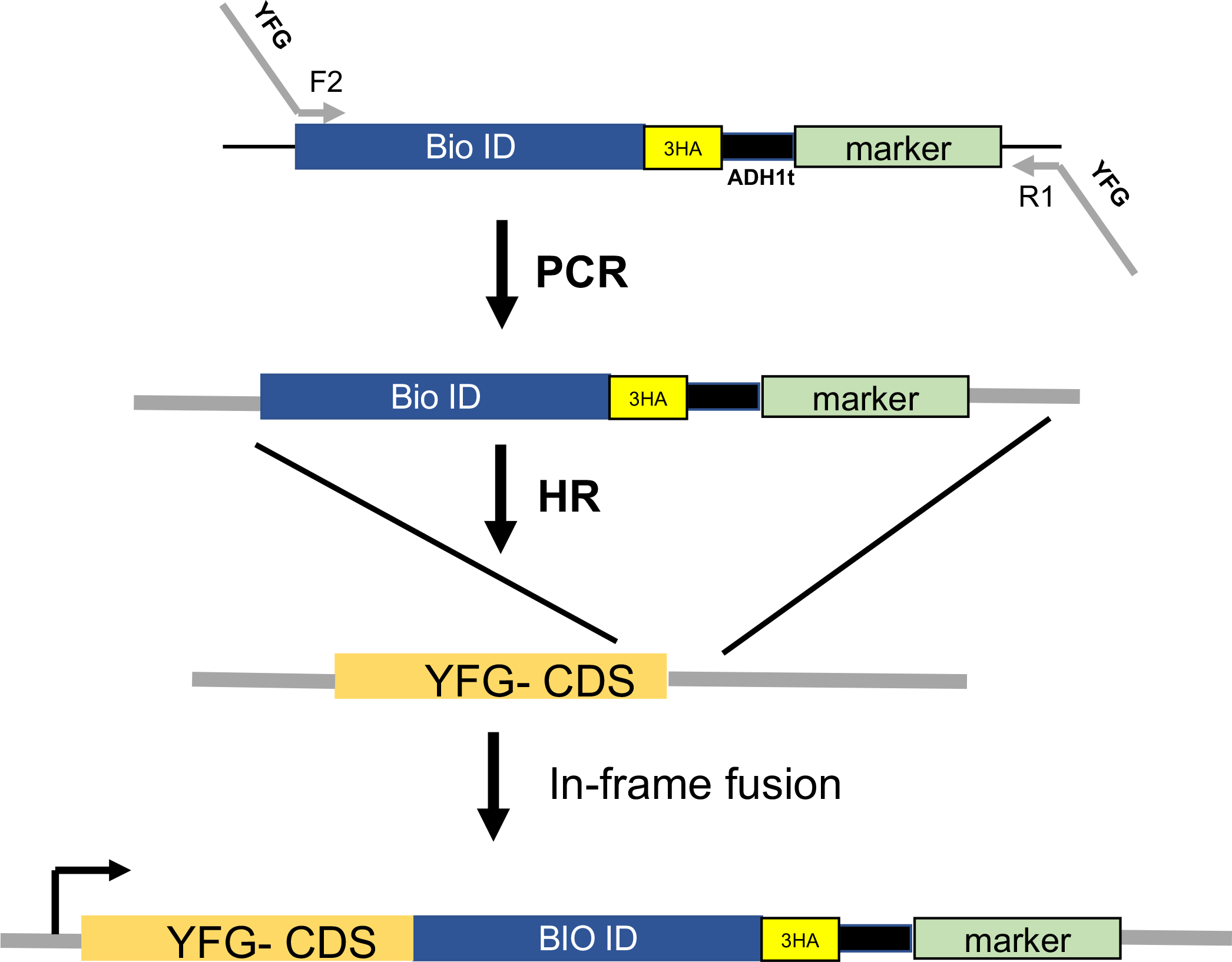
Biotin ligase integration strategy. Plasmids containing BioID enzymes with a triple hemagglutinin epitope tag (3HA) were built into the pF6a vector series. Oligonucleotides containing the F2 and R1 sequences are used to prepare PCR cassettes to integrate the enzyme at the C-terminus of proteins. YFG-your favorite gene; CDS coding sequence.

Next, we grew each strain in synthetic complete medium and added biotin to 10 uM to one of the cultures. Extracts were made, and the levels of the Not4 fusion proteins were detected by western blotting for the HA tag and the amount of biotinylation using streptavidin-horseradish peroxidase (SA-HRP, Figure 2A).

**Figure 2.**
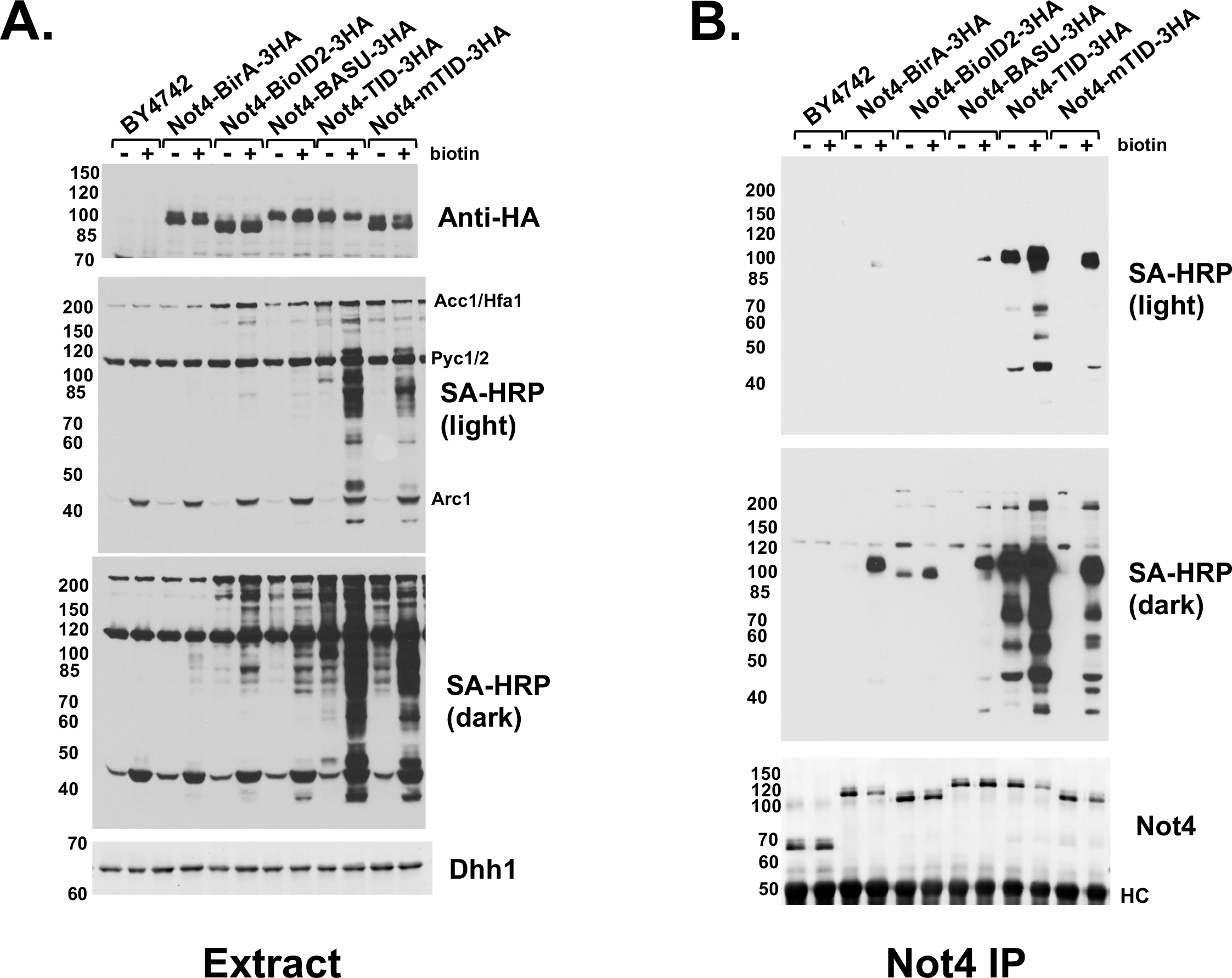
Comparing the activity of biotin ligase enzymes. Strains containing Not4 tagged with the various biotin ligases indicated in the panel and a background control strain BY4741 were grown in synthetic complete media and then treated (+) with or without (-) 10uM biotin for 2 hrs prior to harvest. (A). Western blotting of proteins using anti-HA (12CA5), anti-Dhh1 (loading control) antibodies and streptavidin-HRP (SA-HRP) to detect biotinylated proteins. Two exposures of the SA-HRP blot are presented. (B). Immunoprecipitation of extracts of extracts with anti-Not4 antibodies. Immunoprecipitants were analyzed by SA-HRP blots and western blotting for Not4. HC= heavy chain.

Three major endogenous bands (Acc1/Hfa1, Pyc1/2 and Arc1) were observed in all cells, including in the unmodified strain, BY4742. As noted by others using TurboID in mammalian cells, treating with biotin led to a slight reduction in Not4-TID and Not4- mTID protein (Figure 2A, lanes 3 vs 4, 9 vs 10 and 11 vs 12). The SA-HRP blot revealed that the relative activities of the biotin ligases were TID>mTID>BASU>BioID2>BirA*. Even in the absence of added biotin, the transfer of biotin to proteins was observed with the highly active Not4-TID and Not4-mTID derivatives (Figure 2A, dark). Most yeast strains require biotin, and it is contained in the nitrogen base used to prepare the medium. The Not4-BioID2 fusion also displayed low constitutive biotinylation.

We conducted immunoprecipitation in extracts expressing each of the biotin ligases using a polyclonal antibody to Not4 to confirm the activities of the ligases [15]. As shown in Figure 2B, the relative activities of the biotin ligase enzymes observed in cell extracts was confirmed. Proteins migrating at the expected molecular weights of known Ccr4-Not subunits were observed. Blotting of Not4 protein in the immunoprecipitants confirmed that adding biotin to cells caused reduced protein levels of certain Not4-fusion proteins.

Adding biotin to the cultures grown in a synthetic medium induced the biotinylation of proteins in cells. Since Not4 TurboID (Not4-TID) displayed the highest activity, we characterized this strain further by testing media conditions and biotin induction times. As shown in Figure 3A, cells grown in rich media (YPD) had high, constitutive levels of biotinylation. Adding biotin to YPD did not further increase the levels of biotinylated proteins (data not shown). Treating cells with as little as 1 uM biotin was sufficient for maximal biotin ligase activity. We did not test lower levels of biotin. Our Not4 polyclonal antibody cross-reacts with two weaker bands co-migrating with Not4 (asterisks). We confirmed the signals do not originate from Not4 liberated from the TID tag because they are observed in a not4Δ strain (see [15] and below). To separate the Not4 signal from cross-reacting bands better, we ran the separating gel longer (Figure 3A). As noted in a prior publication, Not4 runs as a doublet resolved on longer gels [13]. Next, we conducted a time course of biotin addition. Biotinylation of proteins was detected by 10 minutes, and the levels started to plateau at 120 minutes. For most of our studies we induce biotinylation for 2 hrs.

**Figure 3.**
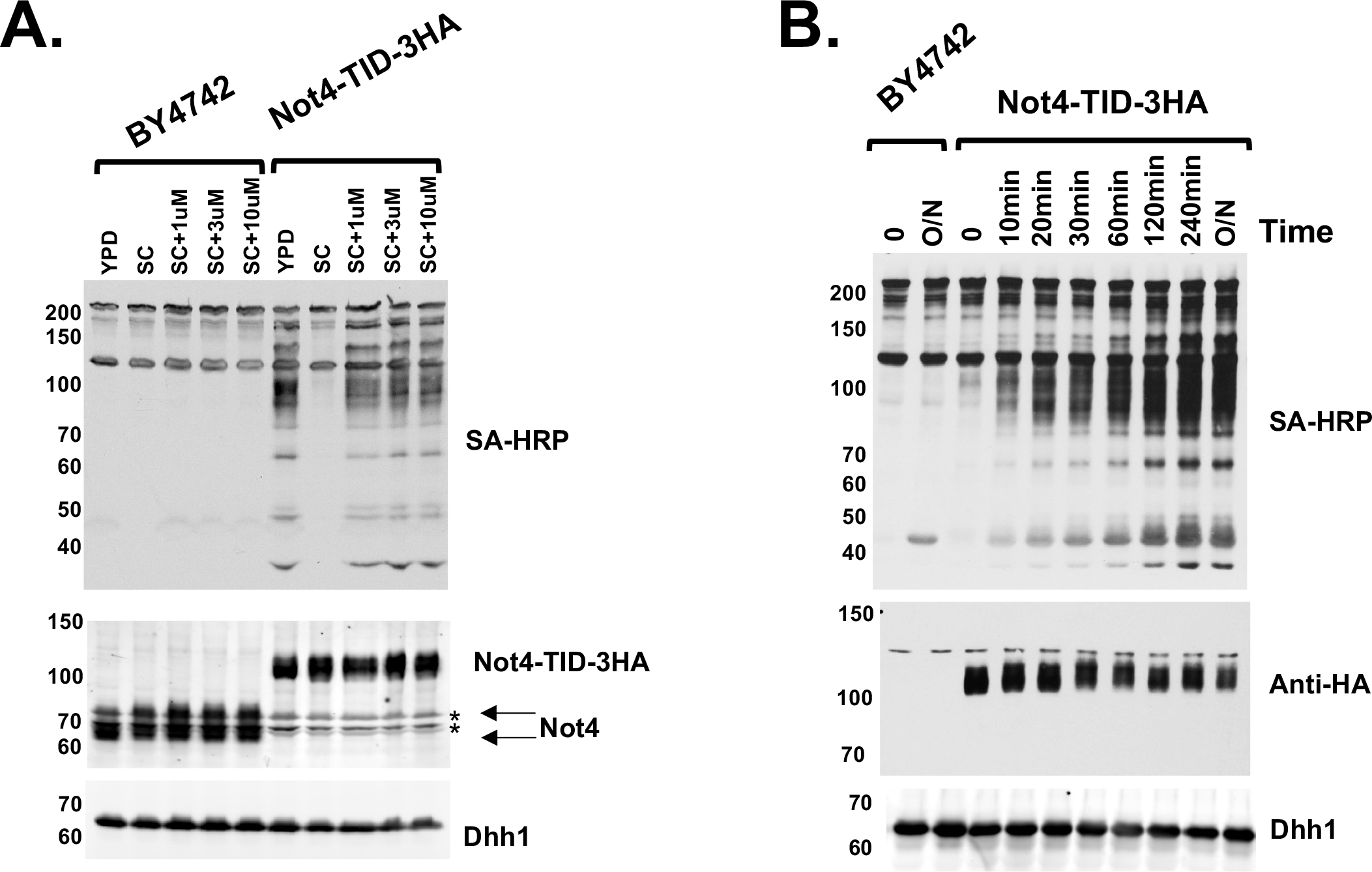
Characterizing the biotin induction conditions for the Not4-TID fusion protein. (A). Extracts of cells were analyzed by SA-HRP blot for biotinylated proteins and western blotting for Not4 and Dhh1 (loading control). The gel was run longer to better separate the Not4 bands from two cross reacting bands (asterisks, see [15] and see Figure 5 for the signal in a *not4Δ* control extract). Cells were grown in rich media (YPD) or synthetic complete (SC) with the concentrations of biotin indicated in the panel. (B). Time course of biotin stimulation. Cells were grown in synthetic complete media and then induced with 1 uM biotin for the times indicated in the panel, O/N= overnight. SA-HRP blots detected biotinylated proteins in cell extracts.

### Chemically induced BioID system

We designed an inducible BioID system based on the rapamycin-induced dimerization of FRB-FKBP (CID-BioID, Figure 4A) [17]. We tagged Not4 at its chromosomal locus on the C-terminus with FRB (Supplemental Table 2). A fusion of TID with FKBP was supplied either on a plasmid copy expressed by yeast promoters of various strengths or by targeting the TID-FKBP fusion downstream of promoters of genes by homologous recombination (Figure 4B and Supplemental Figures 2 and 3).

**Figure 4.**
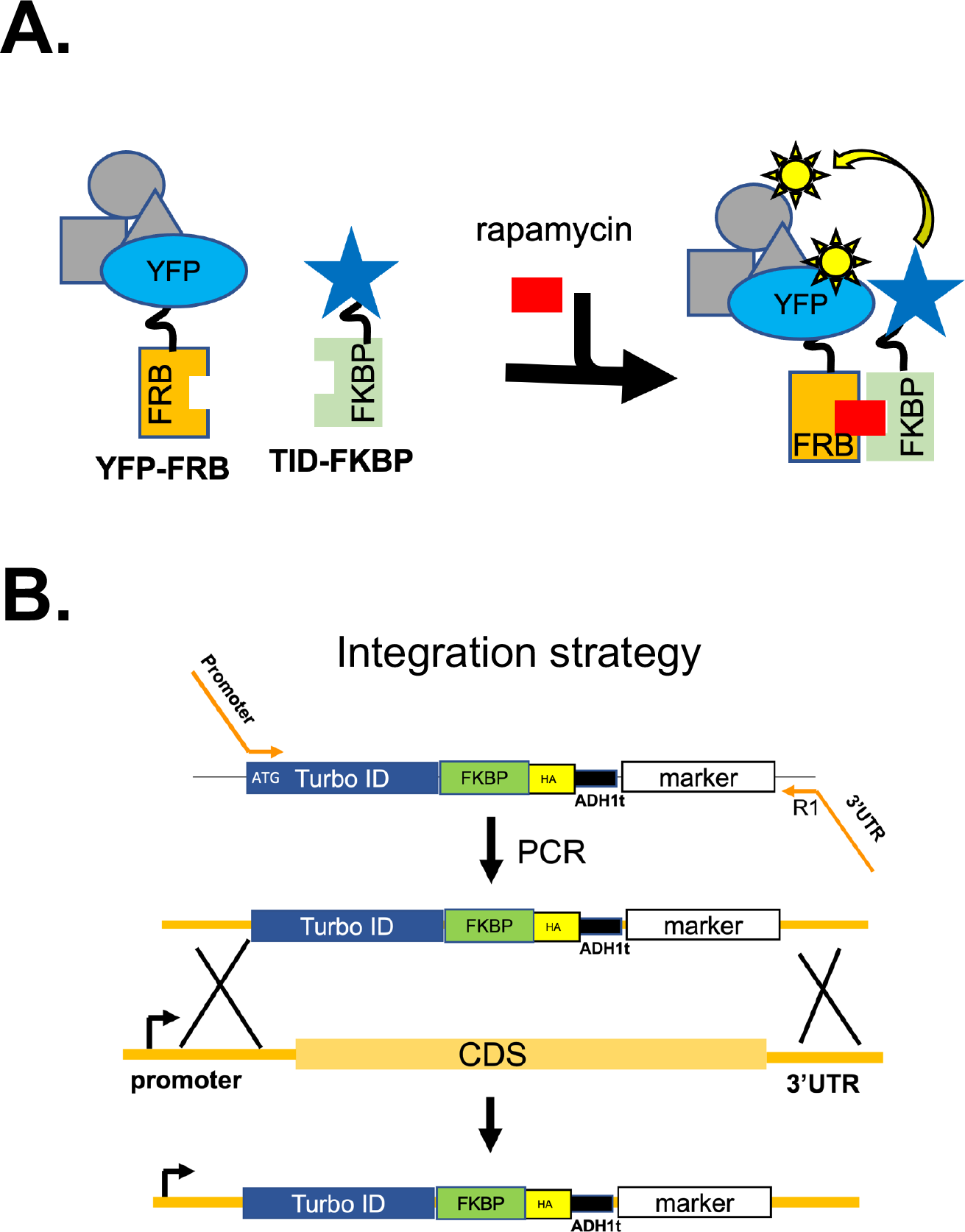
Chemically-induced BioID (CID-BioID) system. (A). Schematic of BioID strategy. (B). integration-based CID-BioID system. A NOT4-FRB strain was transformed with a PCR cassette to integrate TID-FKBP-3HA upstream of yeast promoters.

Integrating the ligase fusion has the advantage of more stable expression, and the TID- FKBP can be controlled by the promoter of virtually any non-essential gene.

First, we tested the CID approach when the FKBP-TID fusion was integrated downstream of different strength promoters, which has the advantage of stable, single-copy expression. The Not4-FRB strain was transformed with a PCR cassette generated from a plasmid to place FKBP-TID under the control of the HIS3 or CUP1 promoter (Figure 4C). We then compared the level of biotinylated proteins from cells treated or not with rapamycin in extracts and Not4 immunoprecipitants (Figure 5).

**Figure 5.**
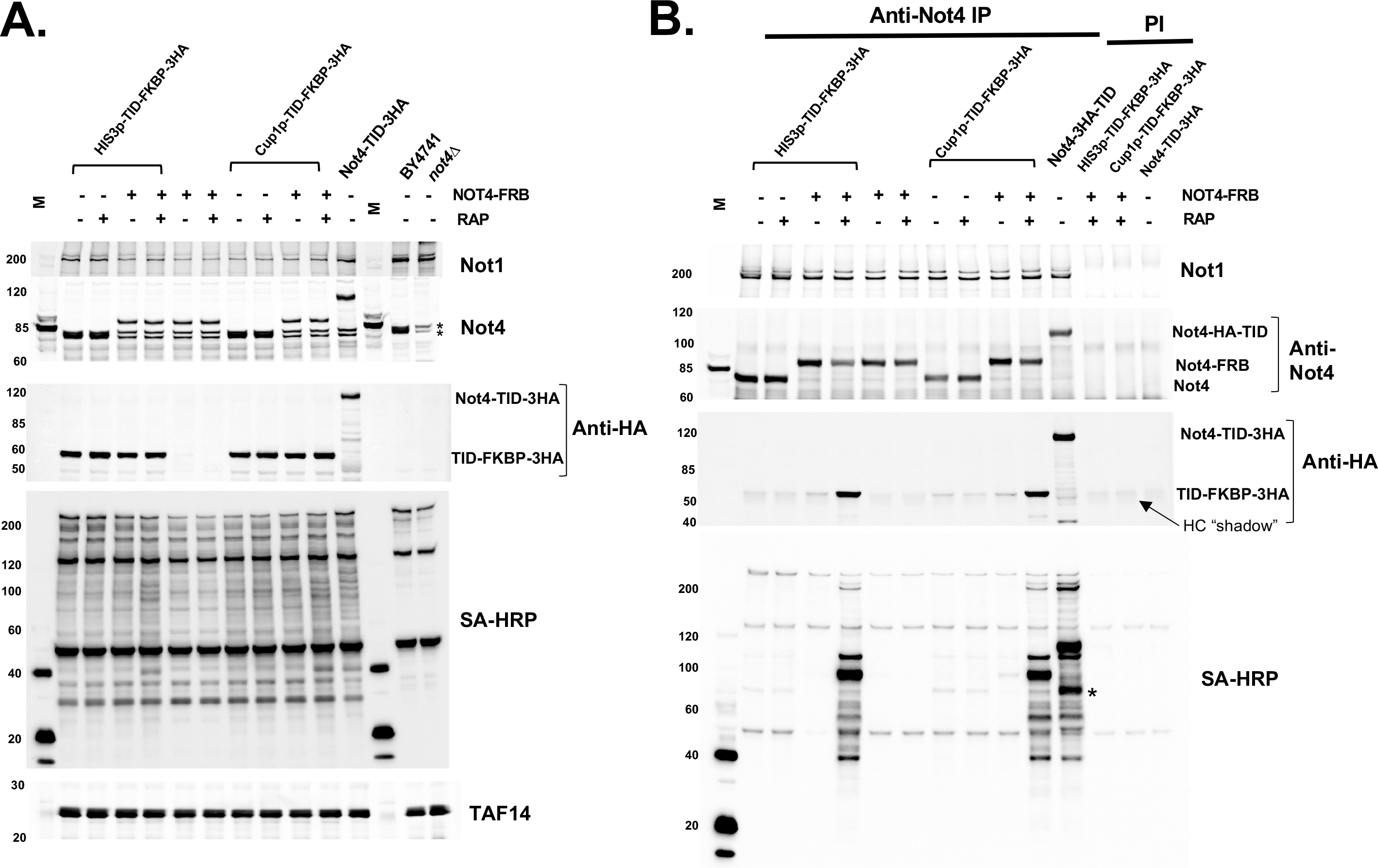
Integrated CID-BioID system. (A). Western and SA-HRP blots of extracts. Cells were grown in synthetic complete media and treated with (+) or (-) rapamycin (RAP) and 1 uM biotin for 2 hrs. Control samples included cells containing only Not4- FRB or no biotin ligase. The lanes labeled NOT4-TID contain samples from a strain where TID was fused to NOT4. Extracts were fractioned on 4-15% gradient gels. The Not4 band co-migrates with the two cross-reacting bands in this gel system. TAF14 is the loading control. “M” is the molecular weight marker lane. (B). Immunoprecipitation of extracts. Extracts were immunoprecipitated with anti-Not4 serum or pre-immune serum (PI, last three lanes). Blotting for Not4 and Not1 determined immunoprecipitation efficiency. The TID-FKBP-3HA fusion protein co-migrates with the heavy chain of the antibody, and weak interaction with the anti-mouse secondary antibody and the rabbit heavy chain produces a “shadow” in the lanes.

The increase in biotinylated proteins in extracts when rapamycin was added was not obvious, but rapamycin-dependent labeling of proteins was observed clearly in the anti-Not4 immunoprecipitants (Figures 5A and B). Comparing the intensity of bands labeled in the immunoprecipitants of the CID-BioID cells with those in cells containing a direct fusion of TID to Not4 (Not4-TID) revealed good but not an exact overlap. One difference is the size of the band corresponding to the auto-biotinylation of the Not4 fusion proteins (identified by overlaying western and biotinylation blots). The other difference is a much stronger labeling of a 70KDa band in the sample where Not4 was fused directly to Not4 (asterisk). Differences in the pattern are unsurprising because inserting the FRB-FKBP proteins between Not4 and the biotin ligase likely changes the distance between the bait and the biotin ligase. Depending on the context, someone can consider this a feature or a drawback of the CID approach. Nonetheless, the chemically-induced system described here works and may be an attractive feature to researchers.

We tested the plasmid system (Supplemental Figures 2 and 3).

Strains containing *NOT4-FRB* fusion transformed with a plasmid expressing TID-FKBP from either the *HIS3* or *CHA1* promoter were used (Figure 4B). The promoter strengths were modest and moderate, respectively. The experiment was performed using time points of 0-120 min of rapamycin treatment (Supplemental Figure 2B).

Biotinylated proteins were detected in extracts and immunoprecipitants using anti-Not4 antibodies. The dimerization of the Not4-FRB and TID-FKBP was confirmed in the IP fraction by blotting for the HA-tag on the TID-FKBP fusion protein. There were some rapamycin-induced bands in the extracts when the TID-FKBP fusion was expressed from the *CHA1* promoter (Supplemental Figure 2C). However, the enrichment of biotinylated proteins in Not4 immunoprecipitants was clearly observed in rapamycin-treated cells (Supplemental Figure 2D). Rapamycin-dependent pulldown of the FKBP-TID fusion protein also was confirmed.

To demonstrate the utility of the CID-BioID system, we applied it to a nuclear protein, TAF1, a subunit of the TFIID complex (Supplemental Figure 3). A strain containing the *TAF1-FRB* fusion was transformed with a plasmid expressing TID-FKBP from the *HIS3*, *CHA1* or *ADH1* promoter. The promoter strengths range from modest, moderate, to high-level expression, respectively. Examining the extract fraction revealed that overexpression FKBP-TID from the strong ADH1 promoter led to spurious biotinylation in the absence of rapamycin. This is likely caused by random collisions between the TID enzyme and cell proteins (and see below). TFIID was immunoprecipitated with anti-TAF1 antibodies. The overexpression of the FKBP-TID from the strong ADH1 promoter led to rapamycin-independent biotinylation of TFIID components (Supplemental Figure 3B). Rapamycin stabilizes the dimerization-competent conformation, and overexpression of the fusion protein induced some dimerization in the absence of the drug. Consistent with that interpretation, blotting for FKBP-TID in the TAF1 IP samples revealed a low level of rapamycin-independent association between TAF1-FRB and FKBP-TID. However, when the fusion protein was moderately expressed, labeling of TAF1-associated proteins required rapamycin (Supplemental Figure 3B). The size of the FKBP-TID protein should allow for its entry into the nucleus. However, we also tested if adding a nuclear localization sequence (NLS) to the FKBP-TID fusion enhanced the efficacy of the assay. We found that the NLS made no difference, however, we did not confirm that the fusion protein accumulated to greater levels in the nucleus when the NLS was added. The NLS-containing versions may be useful to others and will be made available (Table 1).

### Control strains for BioID studies: free biotin ligase expression

BioID studies often compare biotinylated proteins captured from biotin-treated versus untreated cells. This works, but this is not the best control for several reasons. Our experience with expressing biotin ligases from different promoters (Supplemental Figure 3 and data not shown) indicates that non-specific biotinylation of proteins increases with expression levels of the biotin ligase. Even when fused to a bait protein, random collisions with proteins can result in non-target biotinylation. Furthermore, while TurboID has the strongest activity, which is desirable, as shown in Figure 2, and as reported by others, the high activity of TID leads to biotinylation of targets without adding biotin (Figure 2 and [4, 6]). A good control for BioID studies is expressing a “free” biotin ligase to similar levels as the bait-ligase fusion protein. This, combined with isobaric tag (TMT) labeling of samples, provides an excellent strategy to quantify the labeling of target proteins in BioID studies. The plasmid system described above to label proteins with biotin ligases (Figure 1) can be used to produce DNA cassettes to integrate biotin ligases downstream of yeast promoters to express “free” enzyme in cells (Figure 6A). A forward primer containing 42-45 nucleotides homologous to the promoters of genes followed by complementary sequences over the start codon of the ligase and the R1 primer directed to the 3’UTR of the gene is used to generate a PCR product for transformation. The free TID and the bait fusions have the same HA tag, so the investigator can match the expression levels of the free enzyme and the BioID fusion protein. We integrated TID downstream of very strong, moderate, and weak promoters and then compared the free TID protein’s expression level to that of the Not4- and Not1-TID fusions (Figure 6B and C). We also examined the level of biotinylated proteins in the extracts (Figure 6C). As expected, the greater the expression of free TID more protein biotinylation of proteins was observed. Thus, this reinforces the notion that untargeted TID expression leads to the modification of non-target proteins. In this experiment, TID expressed from the HIS3 promoter was the closest match to components of the Ccr4-Not complex and will be used as the control in our BioID studies (Figure 6C). The promoters of CUP1 (copper), CHA1 (hydroxylated amino acids) and MET3 (methionine) have the added feature of expression control by changing media conditions, if necessary. Using a free biotin ligase control strain allows researchers to exploit the benefits of the highly active TurboID enzyme while reducing the drawbacks of promiscuous biotinylation of non-target proteins.

**Figure 6.**
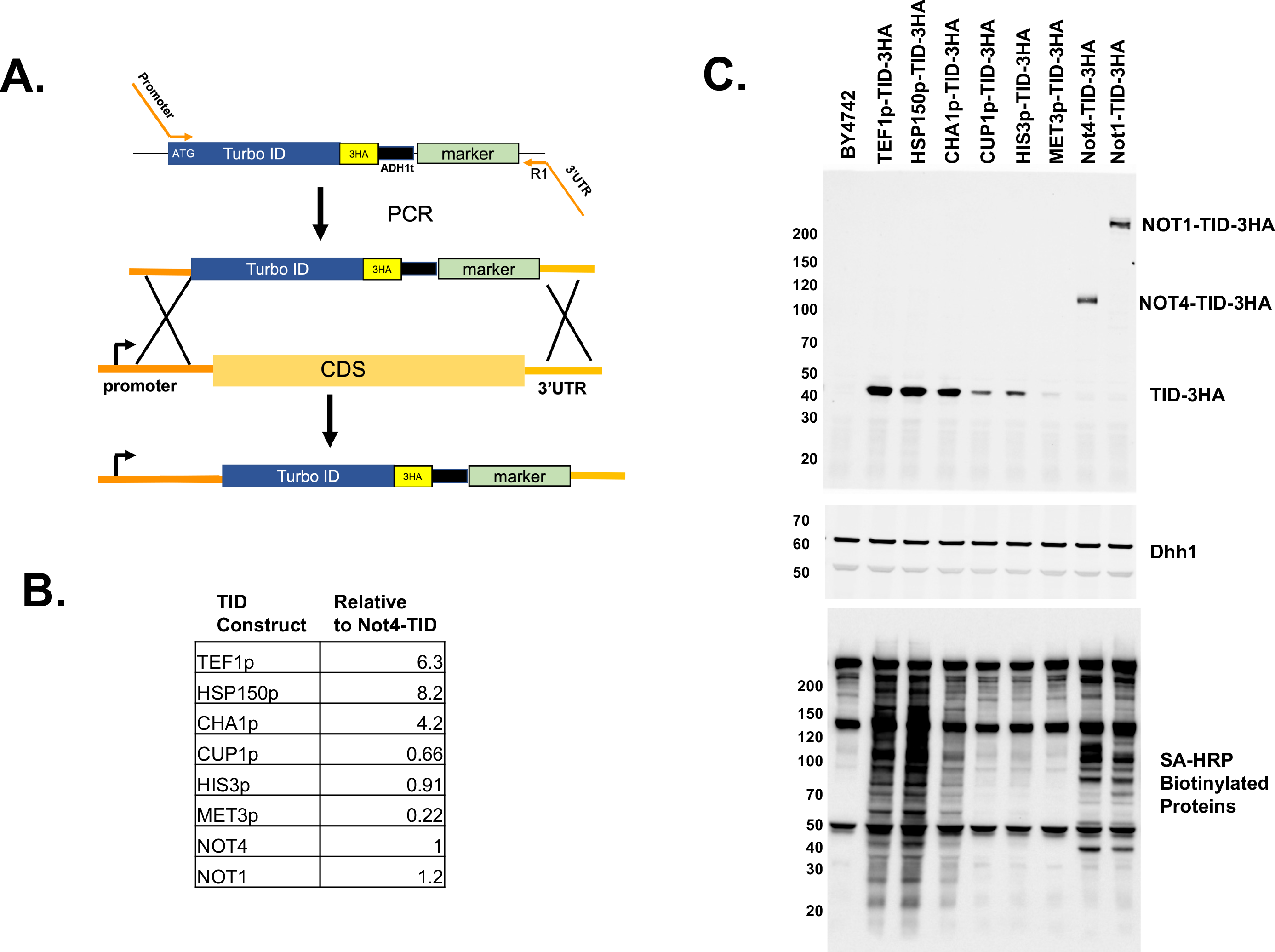
Free enzyme controls for BioID studies. (A). Schematic of preparing strains expressing free TurboID. The same plasmid used to tag genes at the C-terminus of genes is used to produce PCR cassettes, except the forward primer is homologous to the sequences upstream of the start codon of yeast genes and is complementary to the coding sequence of the enzyme coding sequence. (B and C). Blotting and quantification of expression of TurboID (TID) fusion-proteins in extracts. Cells were grown in synthetic media and treated with 1 uM biotin for 2 hrs. (B). The signal from the HA-antibody was normalized to the Dhh1 loading control, and the signal of the Not4-TID-3HA band was set to 1.0. Proteins were separated on 4-15% gradient gels. (C.) The blots used to quantify the signal displayed in panel B.

### Using BioID to map the Interactome of Ccr4-Not

The Ccr4-Not regulates multiple stages of gene expression and ribosome quality control[12–14, 18–21]. The complex was first identified as a regulator of TBP and TFIID and has since been shown to regulate mRNA decay and protein quality control[12, 14, 22]. We used the BioID system described here to explore its interactome. First, we used the Not4-TID strain and free TID expressed from the HIS3 promoter as the control.

After two hours of biotin treatment, extracts were prepared, and biotinylated proteins were isolated under stringent conditions that disrupt non-covalent PPIs (see methods). Proteins captured on streptavidin beads were digested with trypsin, and the purified peptides were labeled with TMT isobaric peptide labeling reagents to quantify the enrichment of proteins in the Not4-TID versus the free TID samples. Each condition was performed in triplicate. The peptide abundances for each replicate showed a strong correlation (R2 =0.9912-0.9992, Supplemental Figure 4). Proteins enriched >1.5 fold and displaying a p<0.05 were identified. The volcano plot in Figure 7A demonstrates that 57 proteins were enriched over the free TID control out of 504 proteins detected in the sample. Eight of the nine subunits of the core Ccr4-Not complex met the criteria, and Not4 displayed the highest enrichment value Figure 7A, black points). Only Not2 was undetected. Not2 is a small protein, and this may explain why it was not identified in the sample or in the samples of the BioID experiments conducted on other subunits of the complex (see below). Dhh1 was also enriched. While not a subunit of the complex, Dhh1 binds to the N-terminal domain of Not1 [13, 23]. Sixteen proteins were underrepresented in the Not4-TID sample. Why proteins would be depleted in this sample is unclear. Fusing a bait to a ligase protein could change its location in the cell, reducing the number of random collisions of the free ligase enzyme with other proteins.

**Figure 7.**
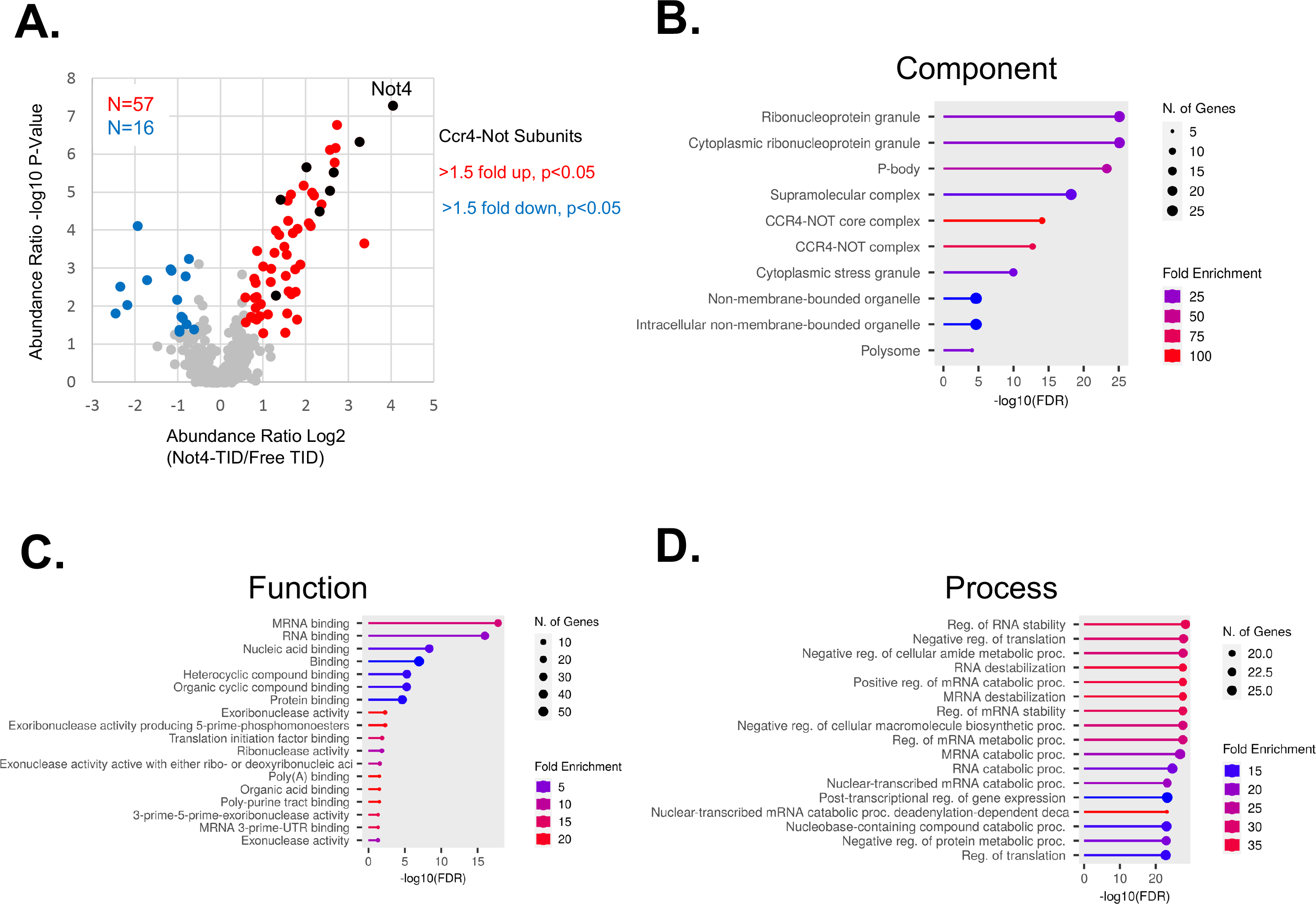
Proximity labeling studies of Not4. Cells were grown in synthetic media and treated with 1 uM biotin for 2 hours before harvesting. Biotinylated proteins were enriched on streptavidin beads and identified by mass spectrometry as described in the methods section. (A). Volcano plot of proteins labeled with Not4-TID versus free TID enzyme expressed from the HIS3 promoter. Proteins enriched >1.5-fold and p<0.05 appear in red, those underrepresented compared to the free TID sample in blue. Ccr4- Not subunits appear as black dots. (B-D) GO terms were identified using the ShinyGo program.

We subjected the enriched proteins to GO term analysis (Figure 7B-D). The most significant terms for cellular components included the Ccr4-Not complex, P-body, cytoplasmic granules, and polysome (Figure 7B). Top function terms included mRNA binding, exonuclease activity, and translation. Process terms were related to mRNA stability (Figure 7D). These GO terms associated with Not4-enriched proteins are entirely consistent with Ccr4-Not’s role in deadenylation and translation control [14, 18–20, 23, 24]. Ccr4-Not functions in the nucleus, including regulating TFIID and transcription elongation [19, 25–28]. Co-immunoprecipitation studies, reconstitution biochemistry experiments and genetic interactions suggest Ccr4-Not binds transcription factors and RNA polymerase II [12, 22, 26–28]. Surprisingly, no subunits of TFIID, transcription elongation factors or RNA polymerase II were identified in the screen. Cell imaging of Ccr4-Not subunits revealed that the majority of Ccr4-Not is cytoplasmic [29, 30]. The proteins associated with the much smaller nuclear fraction of Ccr4-Not may have escaped detection in our screen. Future studies isolating biotinylated proteins from the nuclear fraction may identify Ccr4-Not’s association with nuclear proteins.

Alternatively, the location of the TID enzyme is too far away from the transcription factors interacting with Not4 (although, see below).

Next, we tagged other Ccr4-Not subunits and Dhh1 with Turbo ID (Figure 8A and B). *NOT1* is an essential gene, and deleting *CCR4, CAF1*, or *DHH1* causes slow growth and temperature and hydroxyurea sensitivity [16, 31, 32]. Fusing TID to each protein did not change the growth phenotypes of the cells under the conditions examined, suggesting that, like Not4, these proteins tolerate the fusion with TID (Supplemental Figure 1). Western blotting using anti-HA antibodies revealed that all TID fusion proteins and free TID enzyme were expressed roughly equally (Figure 8B). We used polyclonal antiserum to Not1, Not4, Caf1 and Dhh1 to confirm that the Not1-, Not4- and Dhh1-TID fusion proteins were expressed similarly to the endogenous, untagged protein. The accumulation of the Caf1-TID protein appeared to be lower, but this reduced amount of protein did not alter the growth of the cells (Supplemental Figure 1). Next, we detected the levels of biotinylated proteins in extracts using SA-HRP as a probe (Figure 8A). Fusing TID to Not1, Caf1 and Dhh1 resulted in the appearance of labeling of proteins not observed in the free TID control sample. On the other hand, no obvious new biotinylated proteins were observed in the extract from the Ccr4-TID strain, suggesting low activity of this fusion protein (Figure 8A). We used these strains to conduct BioID studies and compare the results to those obtained for Not4.

**Figure 8.**
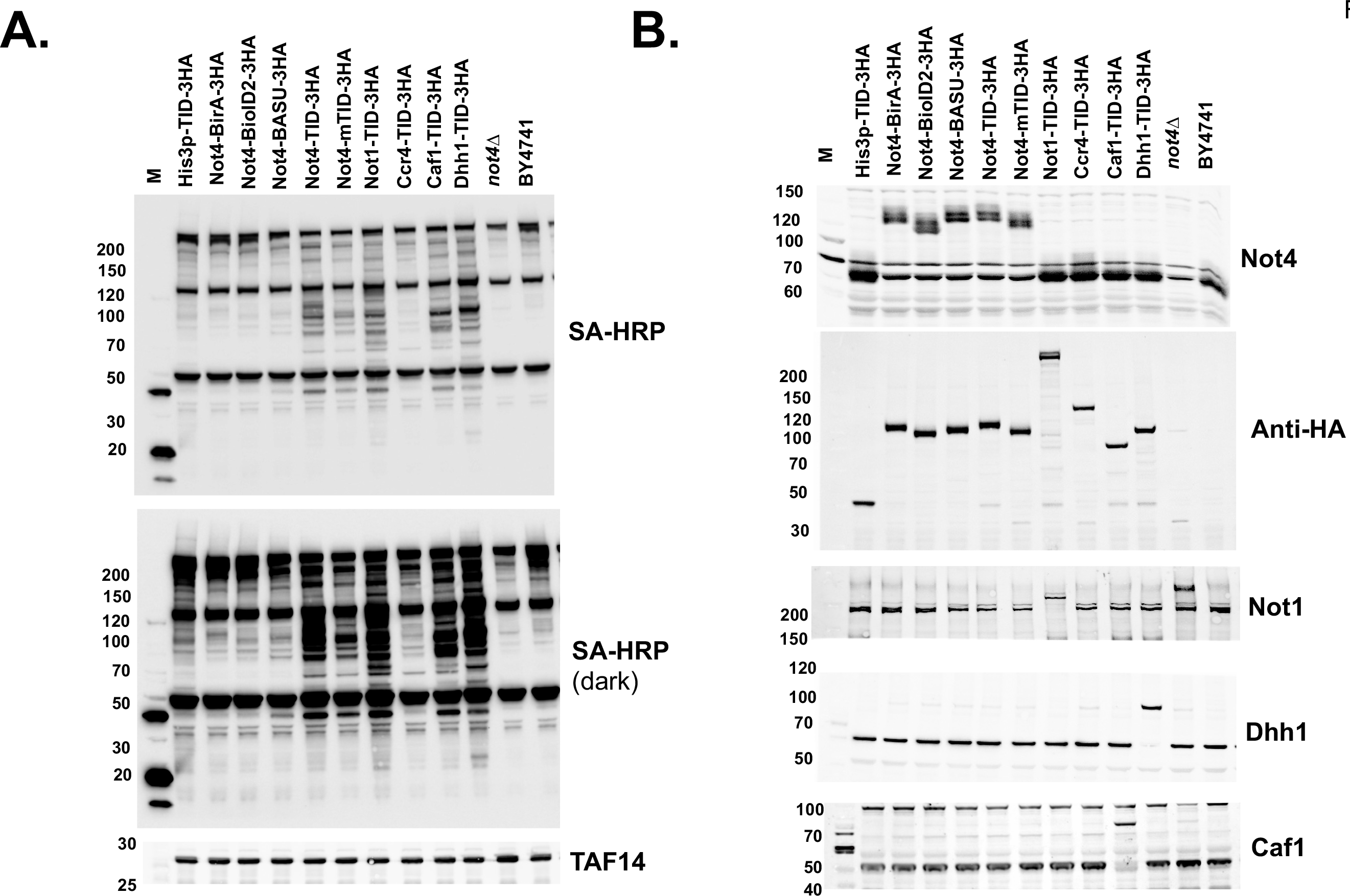
Construction and analysis of BioID strains of multiple Ccr4-Not subunits. Cells were treated as described in Figure 8. (A). Streptavidin-HRP (SA-HRP) detection of biotinylated proteins. M=molecular marker lane. TAF14 is the loading control. Extracts were separated on a 4-15% gradient gel. (B). Western blot of various proteins. The TAF14 blot in “A” comes from the same blot or a duplicate gel run in parallel.

BioID studies were conducted in the same way as for Not4, except they were performed in duplicate. The peptide abundances between the replicates of all samples displayed a high level of correlation (R2 =0.963-0.998, Supplemental Figure 5).

The number of significant “hits” varied among the proteins. Not1 labeled 30 proteins, including seven of the nine subunits of the Ccr4-Not complex (Figure 9A). The data displayed strong overlap with the Not4 data set, where 27 of the 30 Not1 targets were present in the Not4 group (p<3.29E-54) (Figure 10A). BioID on Caf1 identified fewer targets, 14 (Figure 9B), but all 14 overlapped with Not4 (Figure 10B). Five of the proteins labeled by Caf1-TID were in the Ccr4-Not complex, and another subunit fell below the cutoff criteria. The Ccr4-TID data was the weakest of the set. Only seven proteins met the enrichment and statistical cutoffs, but six of the seven were identified in the Not4 collection (Figure 9C and 10C). Even the enrichment values of these seven targets were modest, including that of the bait protein Ccr4 (enrichment 1.65). No other Ccr4-Not subunits were enriched, even though Ccr4 binds directly to Caf1. The low enrichment of targets agrees with the undetectable amount of biotinylation in extracts compared to the other proteins examined here (Figure 8A and B). In fact, there were more depleted proteins, e.g. higher in the free TID control versus the Ccr4-TID sample. This suggests that Ccr4 BioID studies may represent a “fail” (see discussion). Finally, like what was observed for Not4, none of the TID fusion proteins labeled transcription factors (other than Ccr4-Not subunits), and the same GO terms related to cytoplasmic functions were returned for the proteins labeled by each of the subunits (Supplemental Figures 6 and 7).

**Figure 9.**
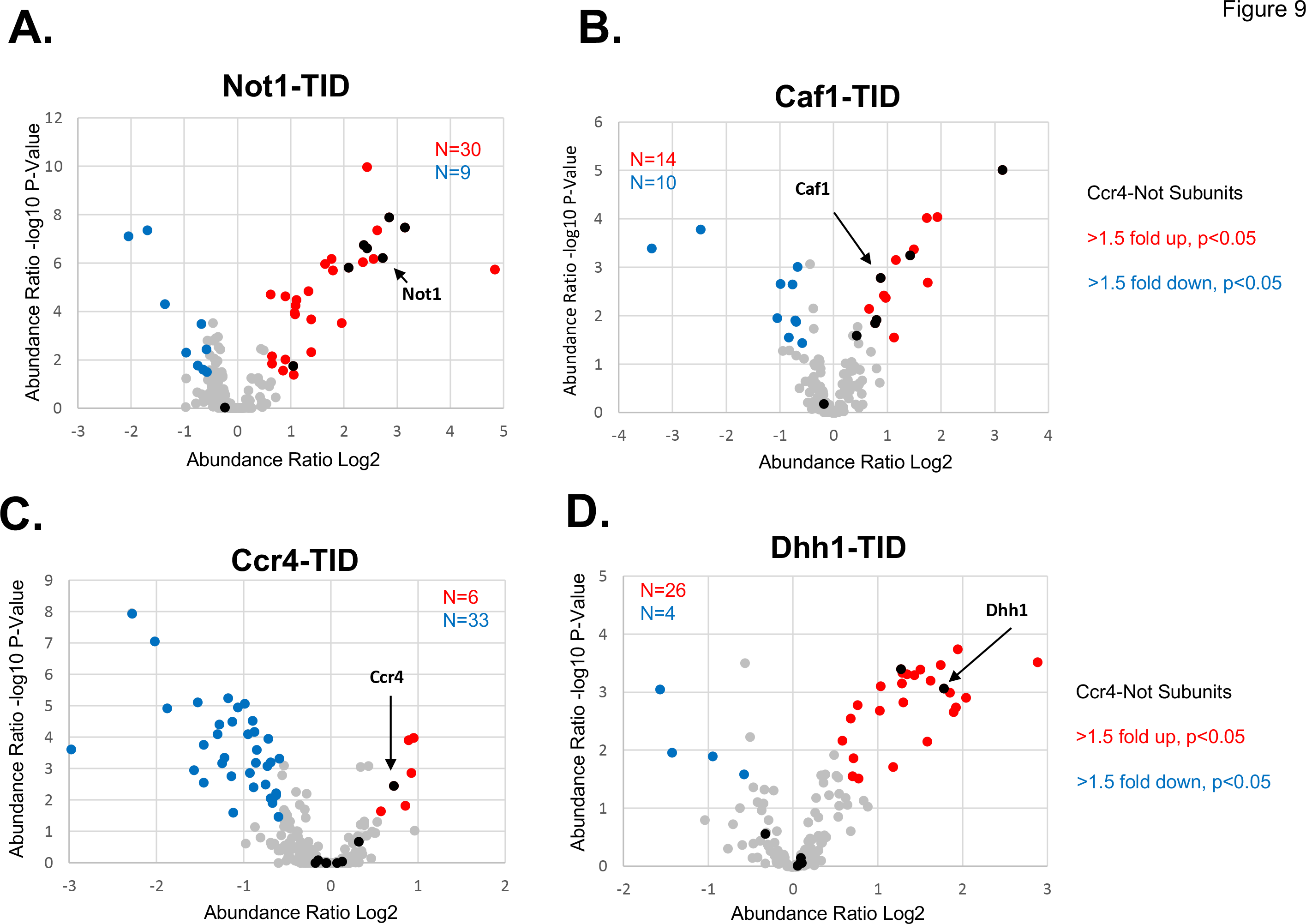
Volcano plots of proximity labeling of multiple Ccr4-Not subunits. (A-D) As in Figure 8. The locations of the bait proteins are indicated in the panels. Free TID expressed from the *HIS3* promoter was used as the control to calculate the enrichment of proteins.

**Figure 10.**
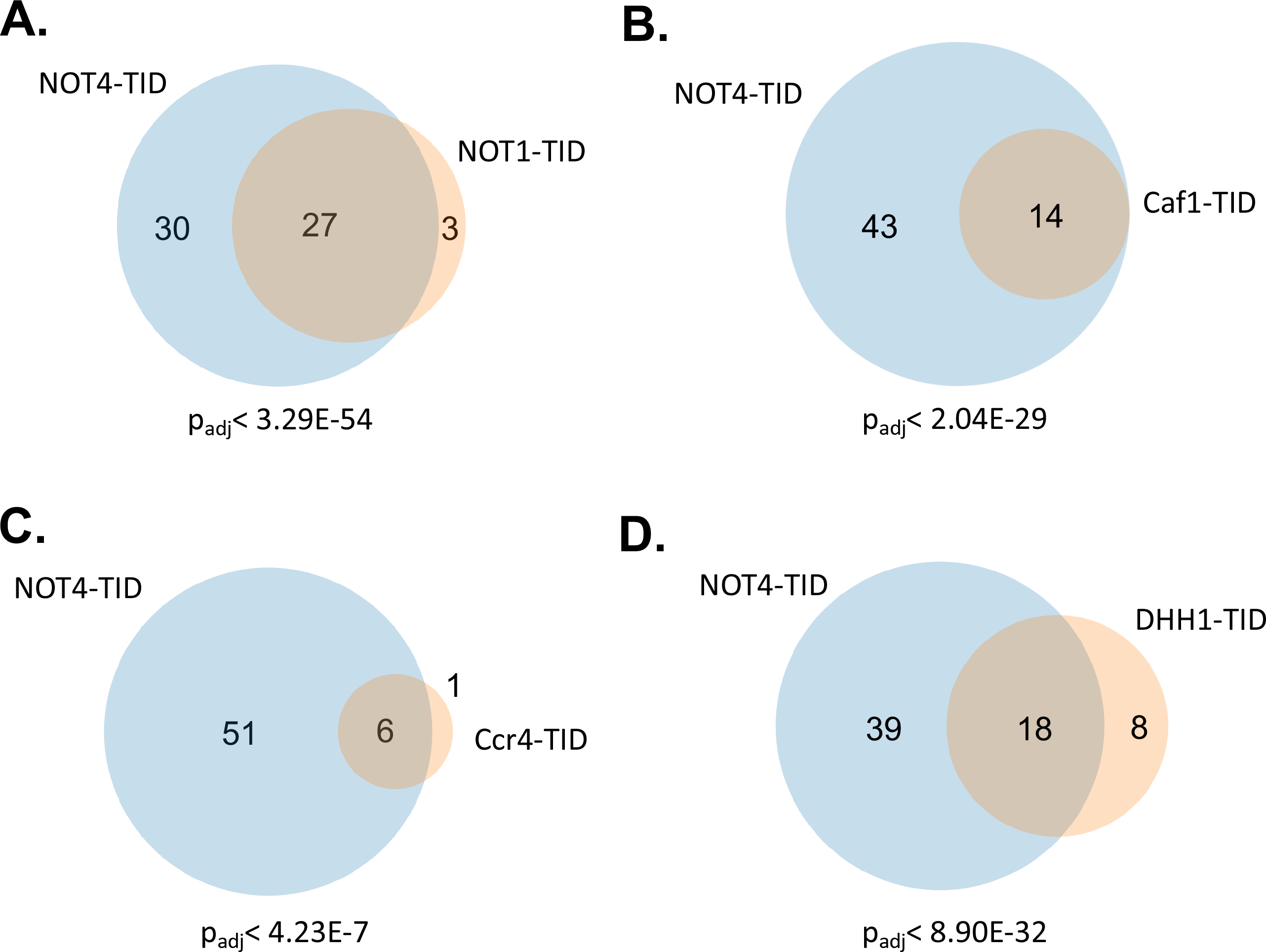
Venn diagrams of the overlap of Not4 interacting proteins and other subunits. The p value under the diagram is the adjusted p value of the overlap.

Dhh1 is not considered a subunit of the complex, but it binds the N-terminus of Not1 [13, 33]. It does not play a role in deadenylation, but like the Ccr4-Not complex, it associates with translating ribosomes and regulates translation and aids in decapping [34–36]. Thus, it has both unique and overlapping functions with Ccr4-Not (Figure 9D). Dhh1 labeled about as many proteins as Not1, but only one Ccr4-Not subunit (Caf40). Nonetheless, 18 of the 26 targets overlapped with that of Not4 (Figure 10D) and 16 of the 26 enriched proteins overlapped with the Not1 BioID data set (Supplemental Figure 8). The GO terms returned from the Dhh1-labeled proteins showed strong overlap with other Ccr4-Not subunits, except for the Ccr4-Not complex component term.

## Discussion

Here, we describe a set of plasmids to conduct BioID in yeast and compare versions of biotin ligase enzymes. We have confirmed that the TurboID versions, TID and mTID, were superior in regards to activity, but they also caused mild instability of the Bait-TID fusion when biotin was added to the culture (Figure 2). It should be noted, however, that the spot growth tests analyzing the phenotypes of all BioID fusion proteins were conducted in rich (YPD) media. Biotin ligase activity was maximal in YPD due to high levels of biotin in the media components (Figure 3). Thus far, we have tagged 12 different proteins with TurboID (TID), and only one strain showed mild growth defects.

Like most tags, adding TID (and other biotin ligases) to the C-terminus of proteins is tolerated. Still, testing the growth properties of every fusion strain under conditions of low (synthetic media) and high activity (YPD) is advisable. Furthermore, whenever antibodies are available, it is a good idea to confirm that the fusion protein’s expression level is comparable to the endogenous protein.

We have found that fusing TurboID to 11 of 12 proteins resulted in robust transfer of biotin to proteins that can easily be detected in extracts and immunoprecipitation studies (Figure 8 and data not shown). This group of proteins includes mRNA decay proteins, translation factors, TFIID and SAGA subunits. Some have not been taken through the mass spectrometry step, but TurboID works well on various proteins.

The one exception that we identified to date is Ccr4. We could not detect biotinylated proteins in extracts, the mass spectrometry results returned few proteins, and the enrichment of biotinylated proteins was low. We only presented data for the Ccr4-TID fusion in this report, but preliminary data suggests mTID, BASU and BirA* fusions of Ccr4 also displayed lower activity compared to the equivalent biotin ligase fusion with Not4 (data not shown). The reason why the Ccr4-fusions “failed” is unclear. The Ccr4- TID strain displayed no mutant phenotypes, including no HU sensitivity, a hallmark phenotype of *CCR4* mutants [16, 32]. Interestingly, BioID on this strain even showed many proteins that displayed higher biotinylation in the free TID samples. One explanation could be that sequences in the C-terminus of Ccr4 may interfere with the folding of the biotin ligase, thus reducing its activity. Placing the biotin ligase on the N-terminus might correct this for Ccr4 and other proteins.

We have also designed a chemically induced BioID system. We demonstrated its effectiveness using two proteins, Not4 and TAF1. We constructed vectors to mix and match the FRB-FKBP pairs (see Table 1). However, we found that the FRB-TID fusion protein is expressed significantly lower than the FKBP-TID version shown here (data not shown). Direct fusion of the biotin ligase to the bait is preferred under most circumstances. However, if a user desires “tighter” control of the induced biotinylation, this system is an attractive alternative. For example, if the goal is mapping lost protein-protein interactions after a stress signal, the CID-BioID would be appropriate. Because proteins display half-lives far greater than typical stress responses, the loss of an interaction would be difficult to detect since the biotinylation of the protein before the stress would remain in the cell for hours. However, if the stress is initiated first and then dimerization and biotinylation are induced, the “lost” proteins would not be biotinylated and detected.

### The proteome of the Ccr4-Not complex

The Ccr4-Not complex was first described as a regulator of TFIID and promoter activity [12, 14]. Multiple studies demonstrated physical and genetic interactions between Ccr4- Not and subunits of the TFIID complex [12, 14, 22]. Purified Ccr4-Not binds RNA polymerase II elongation complexes, and it mediates the DNA damage-dependent degradation of the Rpb1 subunit of polymerase after DNA damage [15, 25–28, 37]. Furthermore, Ccr4-Not co-purifies with TFIIH and crosslinks between subunits of these two complexes have been mapped [38]. We expected to identify proximity interactions with some nuclear proteins. We did not for any of the subunits examined. This was surprising. However, Ccr4-Not is predominantly cytoplasmic, carrying out multiple steps in mRNA decay and translation quality control [12, 14, 19, 29, 30]. The vast majority of the proteins labeled by Ccr4-Not subunit-TID fusion have roles in posttranscriptional gene regulation (Figure 7, and Supplemental Figures 6 and 7). The interactome we mapped represents the cytoplasmic interactions. We hypothesize that the interactions with the smaller fraction of the complex within the nucleus escaped detection.

Future studies that enrich biotin-labeled proteins in the nuclear fraction may identify these interactions. Consistent with what we observe here, BioID conducted on mammalian mRNA regulators using the BirA* biotin ligase also found that human Ccr4- Not predominantly labeled cytoplasmic mRNA regulators, including P-body proteins [39].

All Ccr4-Not subunits, except Ccr4, and Dhh1 strongly labeled Def1 in BioID studies. Def1 was first identified for its role in Rpb1 ubiquitylation and degradation in the nucleus after DNA damage [40]. However, Def1 is predominantly a cytoplasmic protein.

These seemingly contradictory observations were reconciled when it was later found that DNA damage signals stimulate the proteasome-dependent processing of Def1, which allows the N-terminal portion of the protein to accumulate in the nucleus, where it causes Rpb1 ubiquitylation [41]. Likewise, Not4 is required for Rpb1 degradation and is predominantly cytoplasmic [15, 37]. However, we propose that the interaction between Def1 and Ccr4-Not subunits detected by BioID occurs in the cytoplasm. The human homolog of Def1 was recently identified as UBAP2/2L, which is required for Rpb1 ubiquitylation [42]. UBAP2/2L is predominantly cytoplasmic, regulates processing body (P-body) assembly and controls mRNA stability [43, 44].

Proximity labeling studies of RNA binding proteins in human cells found UBAP2/2L labels Ccr4-Not and other P-body components [39]. These observations, and our own, suggest that the Ccr4-Not-Def1 interaction is conserved among eukaryotes.

Considering the many parallels between the function of Def1 and UBAP2/2L, and the conserved protein-protein interactions with cytoplasmic mRNA regulators, it is tempting to speculate that the retention of Def1 in the cytoplasm is not just to keep it from interfering with transcription in the nucleus, but that it has cytoplasmic functions like its human homolog UBAP2/2L.

## Acknowledgements

This research was supported by funds from National Institutes of Health (R35 GM136353 to J.C.R). The mass spectrometry work performed in this work was done by the Indiana University School of Medicine Proteomics Core. Acquisition of the IUSM Proteomics core instrumentation used for this project was provided by the Indiana University Precision Health Initiative. The proteomics work was supported, in part, by the Indiana Clinical and Translational Sciences Institute (funded in part by Award Number UL1TR002529 from the National Institutes of Health, National Center for Advancing Translational Sciences, Clinical and Translational Sciences Award) and, in part, by the IU Simon Comprehensive Cancer Center Support Grant (Award Number P30CA082709 from the National Cancer Institute). Luke Creamer is acknowledged for technical assistance with this work.

