## supplemental figures for "Characterization of BioID tagging systems in budding yeast and exploring the interactome of the Ccr4-Not complex"

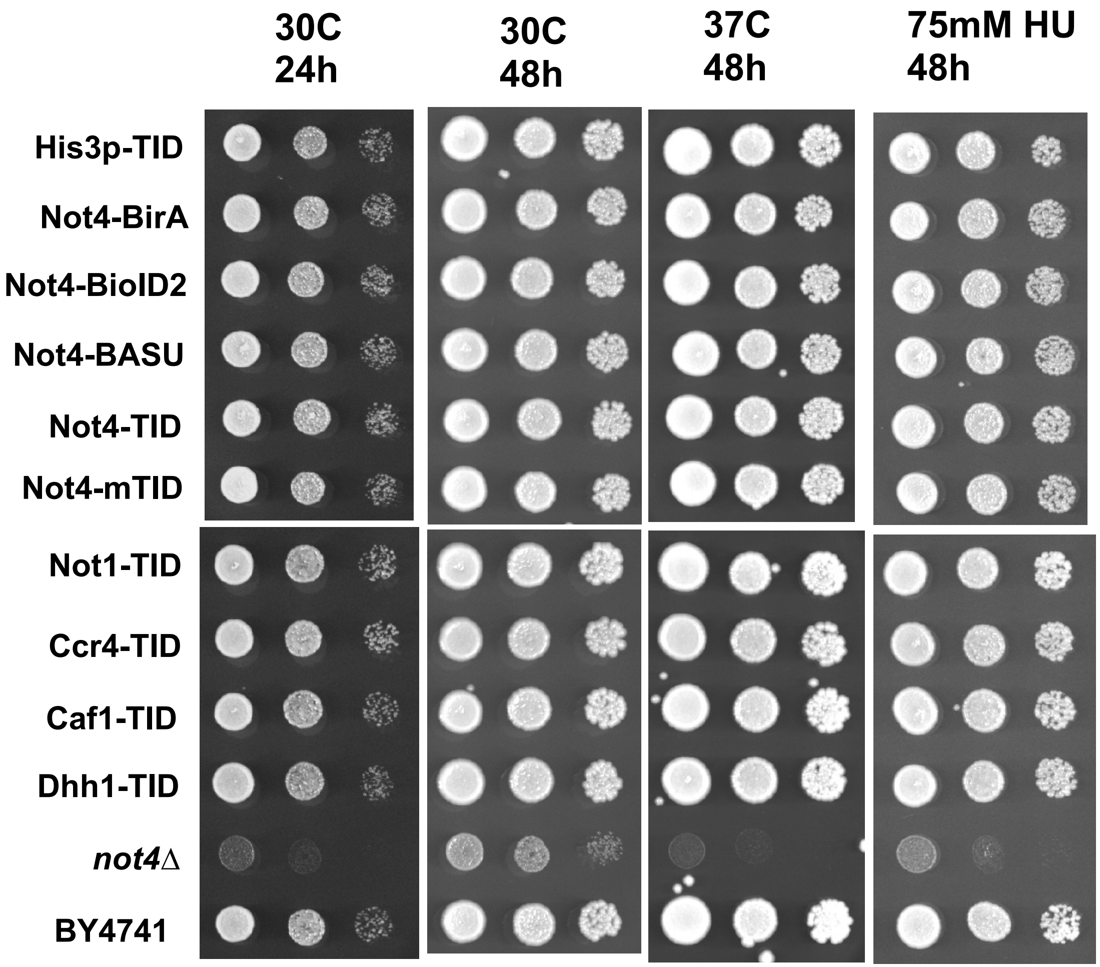
30C
24h
30C
48h
37C
48h
75mM HU
48h
His3p-TID
Not4-BirA
Not4-BioID2
Not4-BASU
Not4-TID
Not4-mTID
Not1-TID
Ccr4-TID
Caf1-TID
Dhh1-TID
*not4Δ*
BY4741

Figure S2

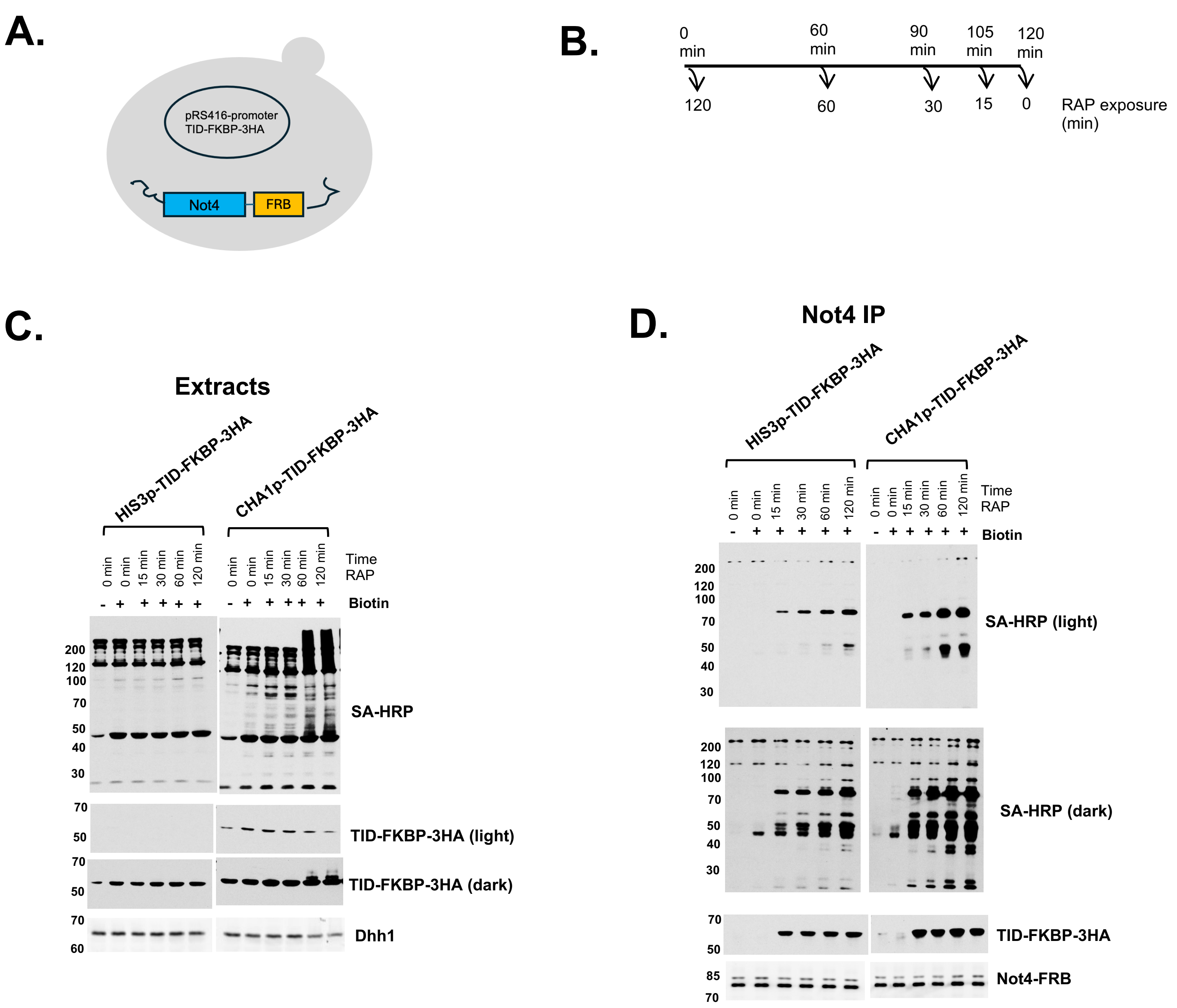
A.
pRS416-promoter
TID-FKBP-3HA
Not4
FRB
B.
0 min
60 min
90 min
105 min
120 min
120
60
30
15
0
RAP exposure (min)
C.
Extracts
HIS3p-TID-FKBP-3HA
CHA1p-TID-FKBP-3HA
0 min
0 min
15 min
30 min
60 min
120 min
Time RAP
Biotin
200
120
100
70
50
40
30
SA-HRP
TID-FKBP-3HA (light)
TID-FKBP-3HA (dark)
60
Dhh1
D.
Not4 IP
SA-HRP (light)
SA-HRP (dark)
TID-FKBP-3HA
85
Not4-FRB

Figure S3

A.

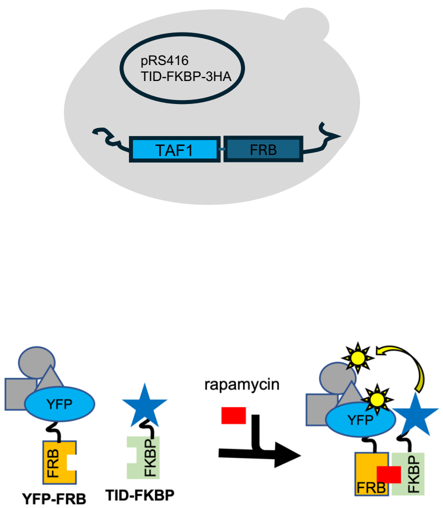
pRS416
TID-FKBP-3HA
TAF1
FRB
YFP
FRB
YFP-FRB
FKBP
TID-FKBP
rapamycin
YFP
FRB
FKBP

B.

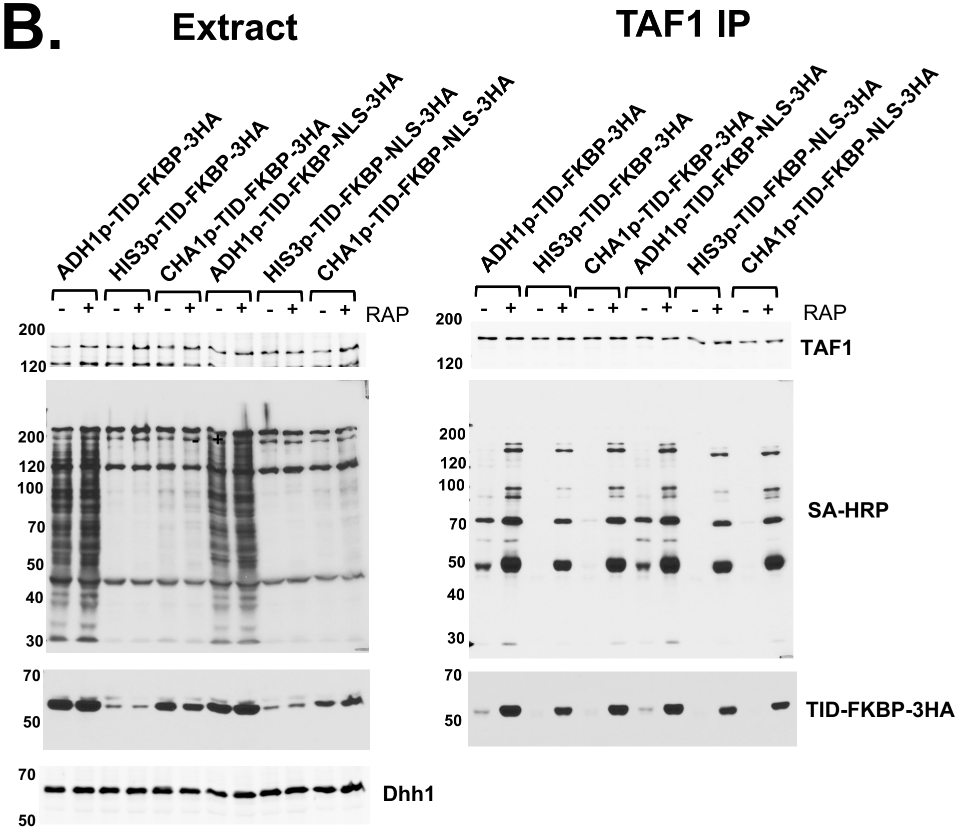
Extract
TAF1 IP
ADH1p-TID-FKBP-3HA
HIS3p-TID-FKBP-3HA
CHA1p-TID-FKBP-3HA
ADH1p-TID-FKBP-NLS-3HA
HIS3p-TID-FKBP-NLS-3HA
CHA1p-TID-FKBP-NLS-3HA
RAP
TAF1
SA-HRP
TID-FKBP-3HA
Dhh1
200
120
100
70
50
40
30

Figure S4

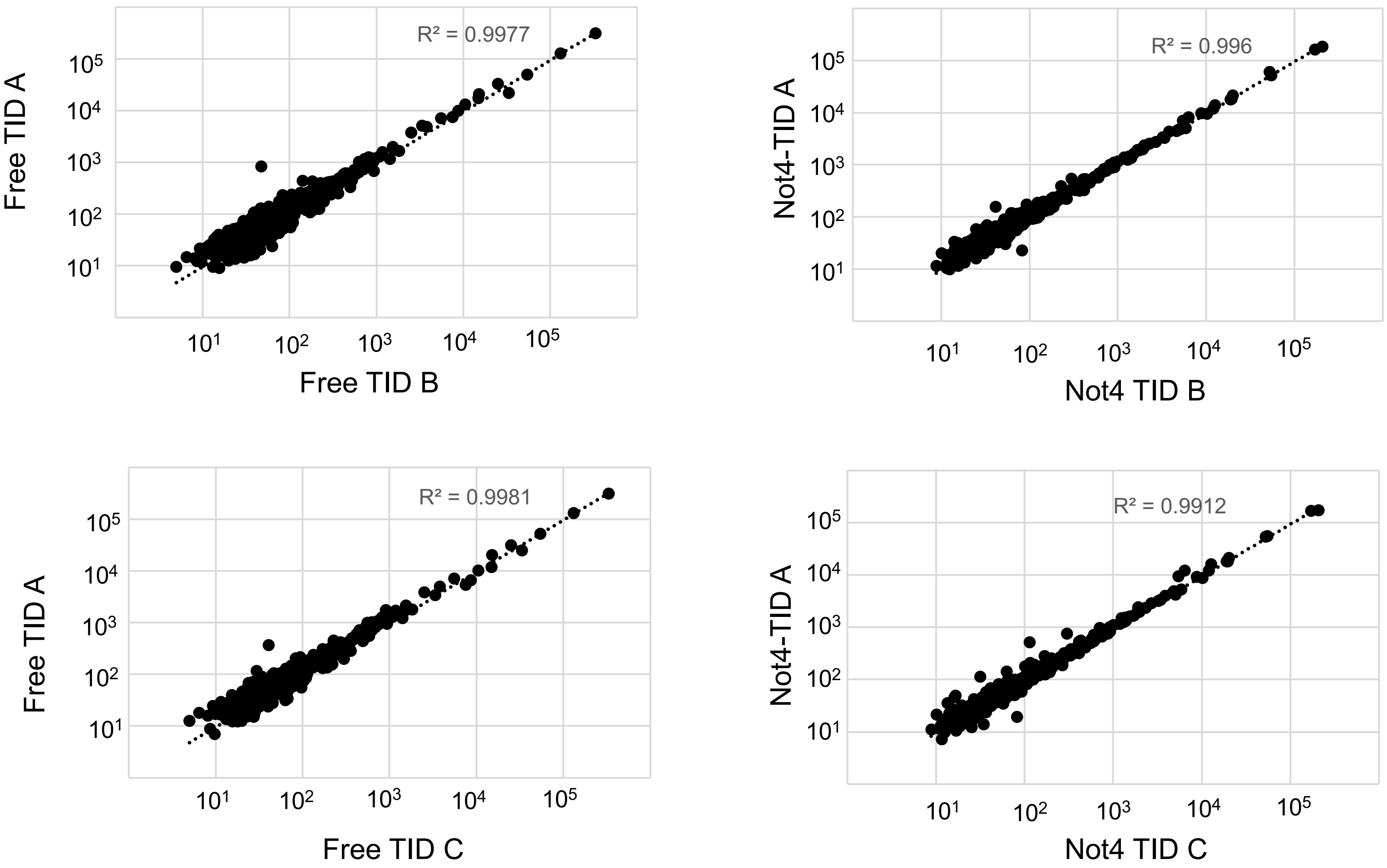

$R^2$ values between replicates

| | Free TID | Not4 TID |
|---|---|---|
| A vs B | 0.9977 | 0.996 |
| A vs C | 0.9981 | 0.9912 |
| B vs C | 0.9992 | 0.9968 |

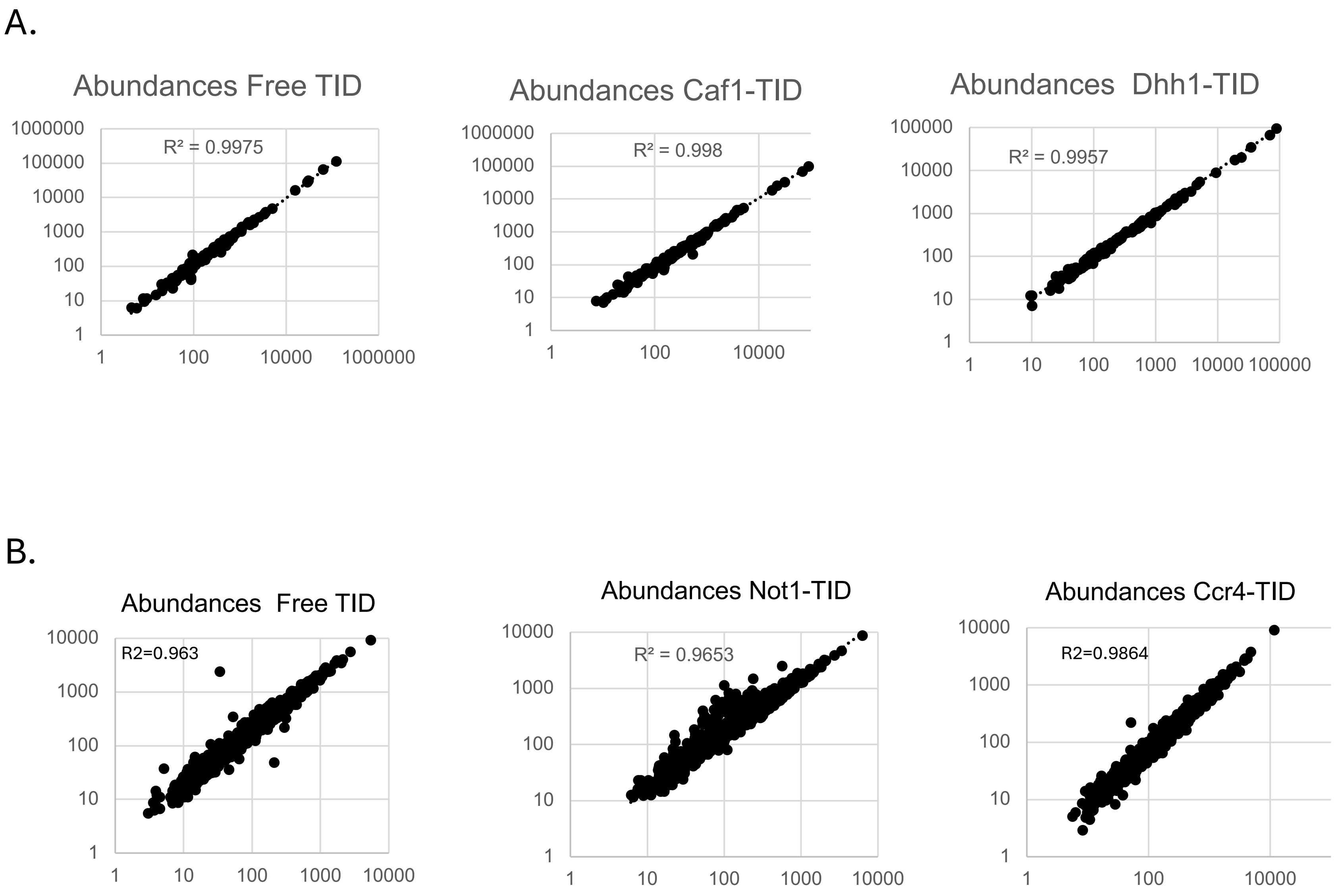
A.
Abundances Free TID
R² = 0.9975
1000000
100000
10000
1000
100
10
1
1
100
10000
1000000
Abundances Caf1-TID
R² = 0.998
1000000
100000
10000
1000
100
10
1
1
100
10000
Abundances Dhh1-TID
R² = 0.9957
100000
10000
1000
100
10
1
1
10
100
1000
10000
100000
B.
Abundances Free TID
R2=0.963
10000
1000
100
10
1
1
10
100
1000
10000
Abundances Not1-TID
R² = 0.9653
10000
1000
100
10
1
1
10
100
1000
10000
Abundances Ccr4-TID
R2=0.9864
10000
1000
100
10
1
1
100
10000

Figure S6

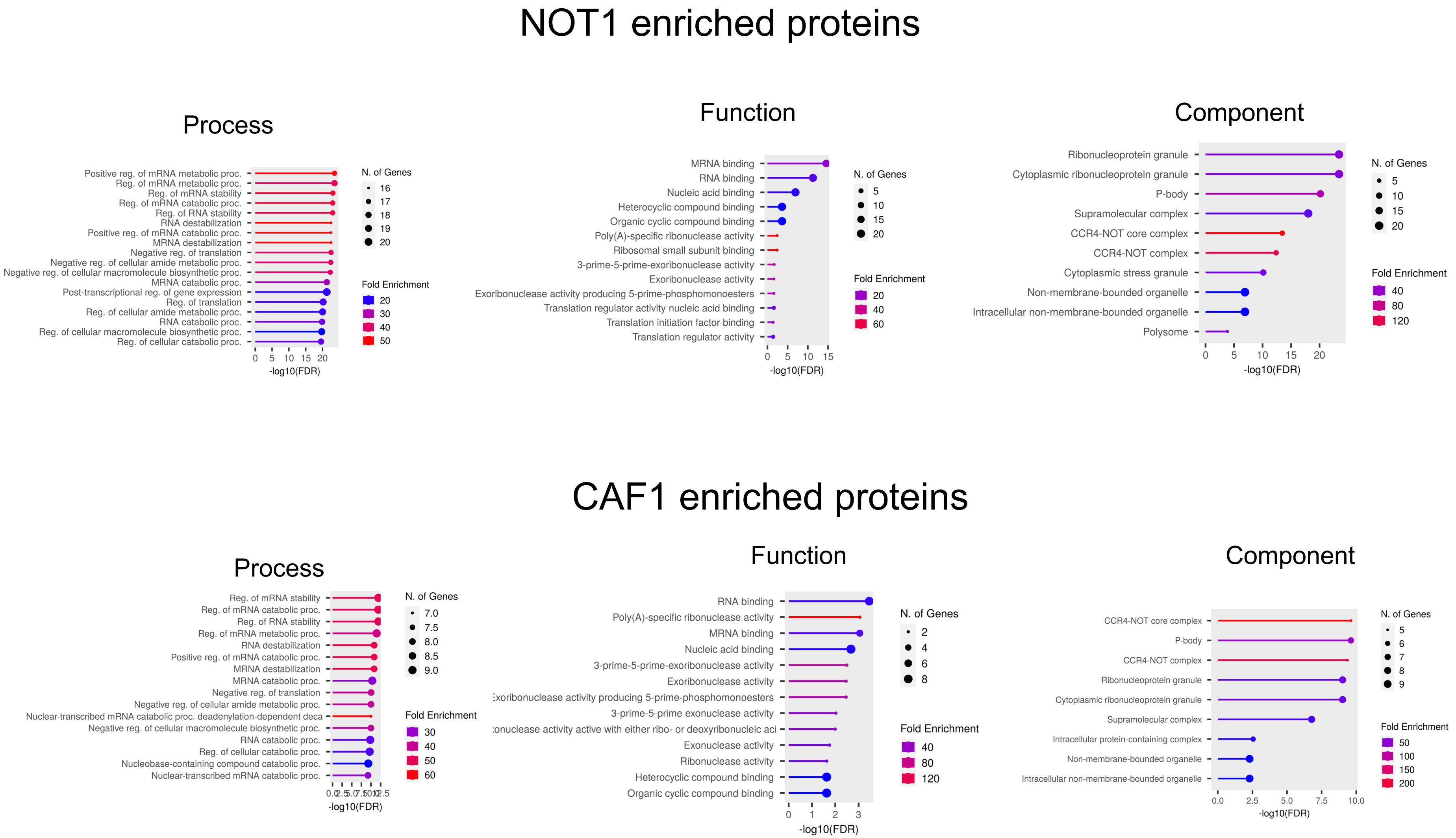
NOT1 enriched proteins
Process
Positive reg. of mRNA metabolic proc.
Reg. of mRNA metabolic proc.
Reg. of mRNA stability
Reg. of mRNA catabolic proc.
Reg. of RNA stability
RNA destabilization
Positive reg. of mRNA catabolic proc.
MRNA destabilization
Negative reg. of translation
Negative reg. of cellular amide metabolic proc.
Negative reg. of cellular macromolecule biosynthetic proc.
MRNA catabolic proc.
Post-transcriptional reg. of gene expression
Reg. of translation
Reg. of cellular amide metabolic proc.
RNA catabolic proc.
Reg. of cellular macromolecule biosynthetic proc.
Reg. of cellular catabolic proc.
0 5 10 15 20
-log10(FDR)
N. of Genes
16
17
18
19
20
Fold Enrichment
20
30
40
50
Function
MRNA binding
RNA binding
Nucleic acid binding
Heterocyclic compound binding
Organic cyclic compound binding
Poly(A)-specific ribonuclease activity
Ribosomal small subunit binding
3-prime-5-prime-exoribonuclease activity
Exoribonuclease activity
Exoribonuclease activity producing 5-prime-phosphomonoesters
Translation regulator activity nucleic acid binding
Translation initiation factor binding
Translation regulator activity
0 5 10 15
-log10(FDR)
N. of Genes
5
10
15
20
Fold Enrichment
20
40
60
Component
Ribonucleoprotein granule
Cytoplasmic ribonucleoprotein granule
P-body
Supramolecular complex
CCR4-NOT core complex
CCR4-NOT complex
Cytoplasmic stress granule
Non-membrane-bounded organelle
Intracellular non-membrane-bounded organelle
Polysome
0 5 10 15 20
-log10(FDR)
N. of Genes
5
10
15
20
Fold Enrichment
40
80
120
CAF1 enriched proteins
Process
Reg. of mRNA stability
Reg. of mRNA catabolic proc.
Reg. of RNA stability
Reg. of mRNA metabolic proc.
RNA destabilization
Positive reg. of mRNA catabolic proc.
MRNA destabilization
MRNA catabolic proc.
Negative reg. of translation
Negative reg. of cellular amide metabolic proc.
Nuclear-transcribed mRNA catabolic proc. deadenylation-dependent deca
Negative reg. of cellular macromolecule biosynthetic proc.
RNA catabolic proc.
Reg. of cellular catabolic proc.
Nucleobase-containing compound catabolic proc.
Nuclear-transcribed mRNA catabolic proc.
0.0 2.5 5.0 7.5 10.0 12.5
-log10(FDR)
N. of Genes
7.0
7.5
8.0
8.5
9.0
Fold Enrichment
30
40
50
60
Function
RNA binding
Poly(A)-specific ribonuclease activity
MRNA binding
Nucleic acid binding
3-prime-5-prime-exoribonuclease activity
Exoribonuclease activity
:xoribonuclease activity producing 5-prime-phosphomonoesters
3-prime-5-prime exonuclease activity
:onuclease activity active with either ribo- or deoxyribonucleic aci
Exonuclease activity
Ribonuclease activity
Heterocyclic compound binding
Organic cyclic compound binding
0 1 2 3
-log10(FDR)
N. of Genes
2
4
6
8
Fold Enrichment
40
80
120
Component
CCR4-NOT core complex
P-body
CCR4-NOT complex
Ribonucleoprotein granule
Cytoplasmic ribonucleoprotein granule
Supramolecular complex
Intracellular protein-containing complex
Non-membrane-bounded organelle
Intracellular non-membrane-bounded organelle
0.0 2.5 5.0 7.5 10.0
-log10(FDR)
N. of Genes
5
6
7
8
9
Fold Enrichment
50
100
150
200

Figure S7

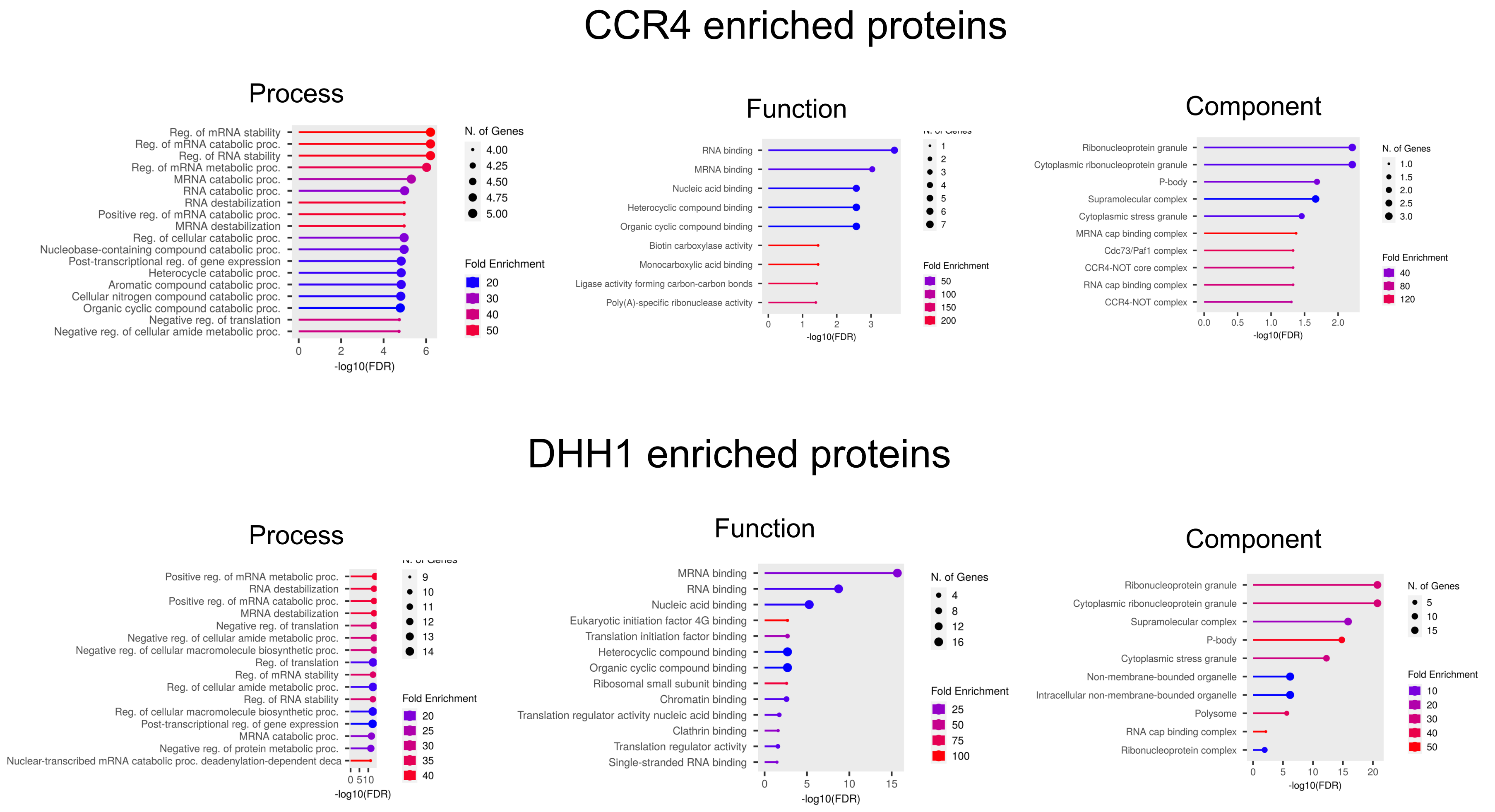
CCR4 enriched proteins
Process
Reg. of mRNA stability
Reg. of mRNA catabolic proc.
Reg. of RNA stability
Reg. of mRNA metabolic proc.
MRNA catabolic proc.
RNA catabolic proc.
RNA destabilization
Positive reg. of mRNA catabolic proc.
MRNA destabilization
Reg. of cellular catabolic proc.
Nucleobase-containing compound catabolic proc.
Post-transcriptional reg. of gene expression
Heterocycle catabolic proc.
Aromatic compound catabolic proc.
Cellular nitrogen compound catabolic proc.
Organic cyclic compound catabolic proc.
Negative reg. of translation
Negative reg. of cellular amide metabolic proc.
-log10(FDR)
N. of Genes
Fold Enrichment
Function
RNA binding
MRNA binding
Nucleic acid binding
Heterocyclic compound binding
Organic cyclic compound binding
Biotin carboxylase activity
Monocarboxylic acid binding
Ligase activity forming carbon-carbon bonds
Poly(A)-specific ribonuclease activity
-log10(FDR)
Fold Enrichment
Component
Ribonucleoprotein granule
Cytoplasmic ribonucleoprotein granule
P-body
Supramolecular complex
Cytoplasmic stress granule
MRNA cap binding complex
Cdc73/Paf1 complex
CCR4-NOT core complex
RNA cap binding complex
CCR4-NOT complex
-log10(FDR)
N. of Genes
Fold Enrichment
DHH1 enriched proteins
Process
Positive reg. of mRNA metabolic proc.
RNA destabilization
Positive reg. of mRNA catabolic proc.
MRNA destabilization
Negative reg. of translation
Negative reg. of cellular amide metabolic proc.
Negative reg. of cellular macromolecule biosynthetic proc.
Reg. of translation
Reg. of mRNA stability
Reg. of cellular amide metabolic proc.
Reg. of RNA stability
Reg. of cellular macromolecule biosynthetic proc.
Post-transcriptional reg. of gene expression
MRNA catabolic proc.
Negative reg. of protein metabolic proc.
Nuclear-transcribed mRNA catabolic proc. deadenylation-dependent deca
-log10(FDR)
Fold Enrichment
Function
MRNA binding
RNA binding
Nucleic acid binding
Eukaryotic initiation factor 4G binding
Translation initiation factor binding
Heterocyclic compound binding
Organic cyclic compound binding
Ribosomal small subunit binding
Chromatin binding
Translation regulator activity nucleic acid binding
Clathrin binding
Translation regulator activity
Single-stranded RNA binding
-log10(FDR)
N. of Genes
Fold Enrichment
Component
Ribonucleoprotein granule
Cytoplasmic ribonucleoprotein granule
Supramolecular complex
P-body
Cytoplasmic stress granule
Non-membrane-bounded organelle
Intracellular non-membrane-bounded organelle
Polysome
RNA cap binding complex
Ribonucleoprotein complex
-log10(FDR)
N. of Genes
Fold Enrichment

Figure S8

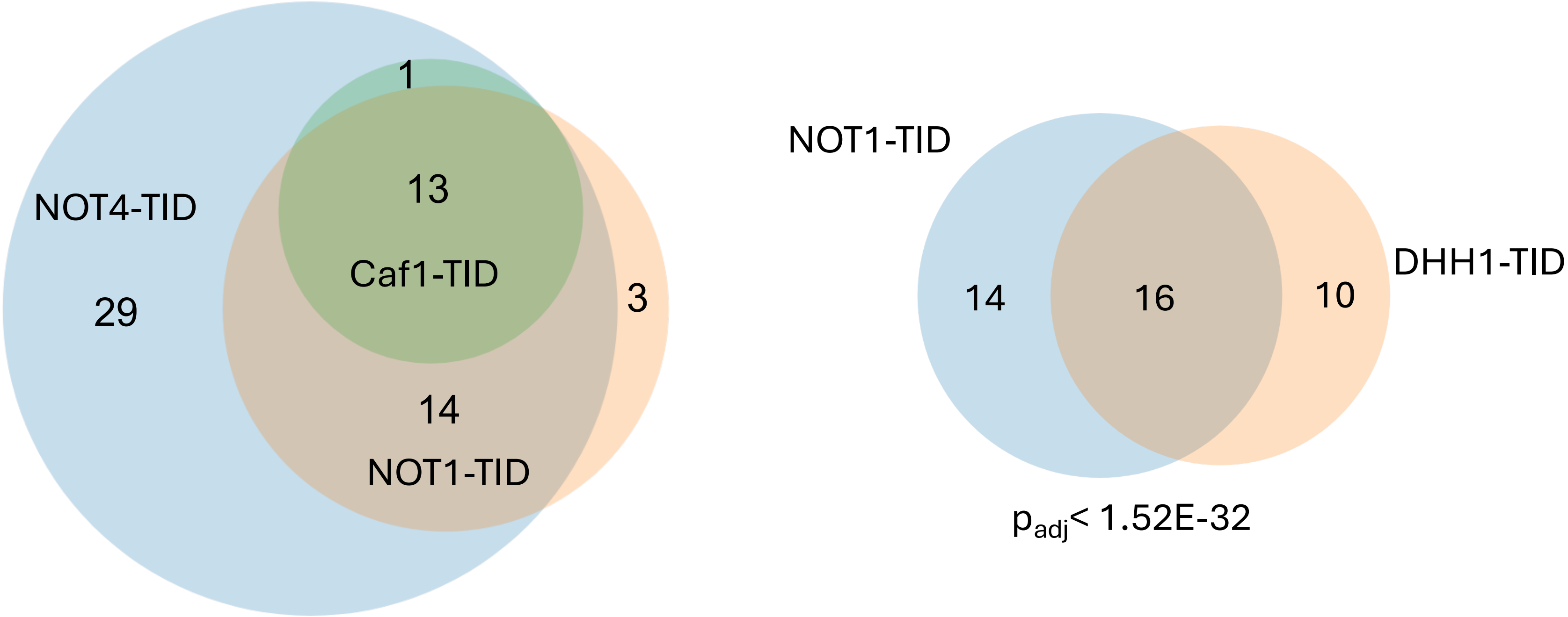
1
13
NOT4-TID
Caf1-TID
29
3
14
NOT1-TID
NOT1-TID
DHH1-TID
14
16
10
$p_{adj}$< 1.52E-32

**Supplemental figure legends**

**Figure S1. Spot test for growth phenotypes of BioID strains.** Cells were grown to saturation in YPD and then diluted to an OD600 1.0, 0.1 and 0.01. Cells were spotted onto YPD or YPD=75 mM hydroxyurea (HU). The plates were incubated for the time indicated above the panels. Not1 is an essential gene, and deletion of Ccr4-Not subunits or *DHH1* causes slow growth, temperature, and HU sensitivity. A *not4*Δ strain is analyzed as a control.

**Figure S2. The plasmid-based CID-BioID system.** (A). Plasmid-based system. A strain expressing Not4 fused to FRB [14]was transformed with a plasmid expressing FKBP-TID-HA3 from different yeast promoters. (B). Schematic of cell treatment regimen. Cells were grown in -uracil media and treated with 1uM biotin and 1 ug/ml rapamycin (RAP) as indicated in panels B and C. All cells were treated with biotin for a total of 2 hrs and the length of rapamycin treatment varied. (C). Blotting of extracts. Strains expressing TID-FKBP-3HA from the *HIS3* and *CHA1* promoter. (C). Immunoprecipitation of extracts using anti-NOT4 antibodies. The gel used to separate Not4 from the heavy chain was run on longer.

**Figure S3. The plasmid-based CID-BioID of TAF1, a nuclear protein.** (A). A strain containing the copy of the *TAF1* gene fused to FRB was transformed with a plasmid with TID-FKBP-3HA expressed from the *HIS3*, *CHA1* and *ADH1* promoters as indicated in panel B. (B). Cells were grown in -uracil media and treated with 1uM biotin and 1 ug/ml rapamycin (RAP) as indicated in panels B and C for 2 hrs. The extract fraction is shown on the left and TAF1 immunoprecipitation is on the right. Overexpressing the FKBP-TID-3HA from the *ADH1* promoter led to rapamycin-independent association of the TAF1-FRB bait and labeling of TAF1-associated proteins.

**Figure S4. Scatter plot of peptide abundances between replicates from the NOT4-TID BioID experiment.**

**Figure S5. Scatter plot of peptide abundances between replicates from the Ccr4-NOT subunit BioID experiment.**

**Figure S6. GO terms of NOT1- and CAF1-enriched proteins.** The collection of enriched protein genes was analyzed by ShinyGo.

**Figure S7. GO terms of Ccr4- and Dhh1-enriched proteins.** The collection of enriched protein genes was analyzed by ShinyGo.

**Figure S8. Venn diagram of the overlap of protein-protein interactions.** (A) Overlap of Not4-, Caf1- and Not1-TID enriched proteins. (B). Overlap between Not1- and Dhh1-TID

### Supplemental Table 1 Yeast strains

| | |
|---|---|
| BY4741 | MAT a his3Δ1; leu2Δ0; met15Δ0; ura3Δ0 |
| BY4742 | MAT alpha his3Δ1; leu2Δ0; lys2Δ0; ura3Δ0 |
| JR1189 | BY4741, not4::KanMx |
| JR1925 | BY4742, NOT4-BirA-3HA::KanMx |
| JR1926 | By4742, NOT4-BioID2-3HA::KanMx |
| JR1927 | BY4742, NOT4-BASU-3HA::KanMx |
| JR1928 | BY4742, CCR4-BirA-3HA::KanMx |
| JR1929 | BY4742, CCR4-BioID2-3HA::KanMx |
| JR1930 | BY4742, CCR4-BASU-3HA::KanMx |
| JR1931 | BY4742, NOT4-TID-3HA::KanMx |
| JR1932 | BY4742, NOT4-mTID-3HA::KanMx |
| JR1933 | BY4742, CCR4-TID-3HA::KanMx |
| JR1934 | BY4742, CCR4-mTID-3HA::KanMx |
| JR2140 | BY4742, CAF1-TID-3HA::KanMx |
| JR2142 | BY4742, DHH1-TID-3HA::KanMx |
| JR1949 | BY4742, TEF1p-TID-3HA::KanMx |
| JR1950 | BY4742, HSP150p-TID-3HA::KanMx |
| JR1951 | BY4742, CHA1p-TID-3HA::KanMx |
| JR1952 | BY4742, CUP1p-TID-3HA::KanMx |
| JR1953 | BY4742, MET3p-TID-3HA::KanMx |
| JR1954 | BY4742, HIS3p-TID-3HA::KanMx |
| JR1963 | BY4742, NOT1-TID-3HA::KanMx |
| JR1918 | MAT alpha tor-1: fpr1:: LoxP-KILEU2-LoxP<br>ade2-11; his3-11,15; leu2-3,112; ura3-1; trp1-1; can1-100 |
| JR1919 | MAT alpha tor-1: frp1::NatMX; NOT4-FRB::KanMX<br>ade2-11; his3-11,15; leu2-3,112; ura3-1; trp1-1; can1-100 |
| JR1937 | MAT alpha tor-1: fpr1:: LoxP-KILEU2-LoxP; TAF1-FRB::KanMx<br>ade2-11; his3-11,15; leu2-3,112; ura3-1; trp1-1; can1-100 |
| JR1966 | MAT alpha tor-1: fpr1:: LoxP-KILEU2-LoxP;<br>HIS3pr-TID-FKBP-3HA::HIS3Mx<br>ade2-11; his3-11,15; leu2-3,112; ura3-1; trp1-1; can1-100 |
| JR1967 | MAT alpha tor-1: fpr1:: LoxP-KILEU2-LoxP<br>HIS3pr-TID-FKBP-NLS-3HA::HIS3Mx<br>ade2-11; his3-11,15; leu2-3,112; ura3-1; trp1-1; can1-100 |
| JR1968 | MAT alpha tor-1: fpr1:: LoxP-KILEU2-LoxP<br>CUP1pr-TID-FKBP-3HA::HIS3Mx<br>ade2-11; his3-11,15; leu2-3,112; ura3-1; trp1-1; can1-100 |
| JR1969 | MAT alpha tor-1: fpr1:: LoxP-KILEU2-LoxP<br>CUP1pr-TID-FKBP-NLS-3HA::HIS3Mx<br>ade2-11; his3-11,15; leu2-3,112; ura3-1; trp1-1; can1-100 |
| JR1970 | MAT alpha tor-1: fpr1:: LoxP-KILEU2-LoxP;<br>HIS3pr-TID-FRB-3HA::KanMx<br>ade2-11; his3-11,15; leu2-3,112; ura3-1; trp1-1; can1-100 |
| JR1971 | MAT alpha tor-1: fpr1:: LoxP-KILEU2-LoxP<br>HIS3pr-TID-FRB-NLS-3HA::KanMx<br>ade2-11; his3-11,15; leu2-3,112; ura3-1; trp1-1; can1-100 |
| JR1972 | MAT alpha tor-1: fpr1:: LoxP-KILEU2-LoxP<br>CUP1pr-TID-FRB-3HA::KanMx |

| | |
|---|---|
| | ade2-11; his3-11,15; leu2-3,112; ura3-1; trp1-1; can1-100 |
| JR1973 | MAT alpha tor-1: fpr1:: LoxP-KILEU2-LoxP<br>CUP1pr-TID-FRB-NLS-3HA::KanMx<br>ade2-11; his3-11,15; leu2-3,112; ura3-1; trp1-1; can1-100 |
| JR1978 | MAT alpha tor-1: fpr1:: LoxP-KILEU2-LoxP; NOT4-FRB-KanMx<br>HIS3pr-TID-FKBP-3HA::KanMx<br>ade2-11; his3-11,15; leu2-3,112; ura3-1; trp1-1; can1-100 |
| JR1979 | MAT alpha tor-1: fpr1:: LoxP-KILEU2-LoxP; NOT4-FRB-KanMx<br>CUP1pr-TID-FKBP-3HA::HIS3Mx<br>ade2-11; his3-11,15; leu2-3,112; ura3-1; trp1-1; can1-100 |
